# The proximal proteome of FLOWERING LOCUS T LIKE 1 during rice panicle development suggests cell-to-cell mobility features

**DOI:** 10.1101/2025.07.06.662997

**Authors:** Daniele Chirivì, Giulia Ave Bono, Jeroen de Keijzer, Ludovico Dreni, Franco Faoro, Francesca Giaume, Cristina Ferrándiz, Fabio Fornara, Camilla Betti

## Abstract

Flowering is promoted by perception of favorable environmental stimuli. In rice, exposure of leaves to short days induces the differentiation of a branched inflorescence called panicle at the shoot apical meristem (SAM). Systemic communication from the leaves to the SAM is mediated by members of the phosphatidylethanolamine-binding protein (PEBP) family, including HEADING DATE 3a (Hd3a) and RICE FLOWERING LOCUS T 1 (RFT1), commonly referred to as ‘florigens’. Florigens are ∼20kDa proteins translated in leaf companion cells and loaded into the phloematic stream, through which they reach the SAM. Once in meristematic cells, they are translocated to the nucleus, where they form higher-order protein complexes that include transcription factors and drive transcriptional reprogramming of meristematic cells. The activity of Hd3a and RFT1 at the SAM is partly mediated by *FLOWERING LOCUS T LIKE 1* (*FTL1*) encoding a florigen-like protein required to accelerate flowering and establish the branching pattern of the panicle. The photoperiodic regulatory network is largely based on protein-protein interactions (PPIs), that take place in different tissues and in response to environmental variation. Capturing the full extent of these interactions for a specific protein and *in vivo* is a challenging task. Here, we developed a protocol of Proximity Labeling (PL) to study the interactome of FTL1 in developing panicles. We show that FTL1 associates with, or is proximal to, several proteins implicated in vesicular trafficking, microtubule binding and transcriptional regulation. Using a transient system, we demonstrate that FTL1 can actively move between cells, similarly to Hd3a and RFT1. This study establishes a protocol to implement PL for the study of a dynamic process during rice reproductive development, identifies interactors of FTL1, and suggests novel features possibly linked to its function.

## INTRODUCTION

In plants, the control of flowering time underlies the important change of switching to sexual reproduction. Environmental and autonomous information induce the floral transition through a strict series of molecular steps, ultimately leading to the differentiation of the shoot apical meristem (SAM) into the reproductive organs (Chirivì & Betti, 2023; Freytes et al., 2021; Kean-Galeno et al., 2024; Tsuji & Sato, 2024). In *Oryza sativa*, the floral transition prompts the SAM to transform into an inflorescence meristem (IM), which sequentially generates higher order meristems until the floral organs develop. These include primary and secondary branch meristems (PBM and SBM), spikelet meristems (SM), which produce florets and glumes, and floret meristems (FM), which form the flowers. The resulting inflorescence structure is called panicle, and the timing of meristematic transitions determines its final architecture, including the number of branches and flowers (G. Li et al., 2021; Vicentini et al., 2023).

All biological processes, such as the transition from vegetative to reproductive growth in plants, rely on the establishment of protein-protein interactions (PPIs), which may induce changes in physical structures or activation/deactivation of protein functions. For example, transcription of genes required for the transition to flowering results from multiple interactions among regulatory proteins and often involves the integration of environmental cues into complex gene regulatory networks (GRNs) (Y.-C. Wang & Chen, 2010). The derived network of PPIs is quite complex and has a high plasticity to quickly respond to different inputs and is usually referred to as “interactome”.

In rice, the transition to the reproductive phase is accelerated when day length (photoperiod) falls below a critical threshold (Goretti et al., 2017; Itoh et al., 2010). Short day (SD) conditions are perceived in leaves by photoreceptors, including phytochromes (PHY) and cryptochromes which integrate light quantity/quality information into GRN. The control of PPIs is necessary to confer a photoperiodic response. For example, PHYA and PHYB interact with GRAIN NUMBER, PLANT HEIGHT AND HEADING DATE 7 (Ghd7), to reduce its accumulation under SD (Zheng et al., 2019). Ghd7 is a long day (LD) flowering repressor belonging to the CCT family of transcription factors and binds the DNA by forming higher-order complexes with NUCLEAR FACTOR Y (NF-Y) B and C subunits (Chaves-Sanjuan et al., 2021; Goretti et al., 2017; Shen et al., 2020). Similarly, HEADING DATE 1 (Hd1) and PSEUDO RESPONSE REGULATOR 37 (PRR37), also encoding CCT family members, can establish trimeric interactions with NF-YB/C (Goretti et al., 2017; Shen et al., 2020). Hd1 and the rice-specific B-type response regulator EARLY HEADING DATE 1 (Ehd1) act at the core of the photoperiodic flowering network (Biancucci et al., 2025; Doi et al., 2004; Yano et al., 2000). Under SD, they are necessary to promote the transcription of *HEADING DATE 3a* (*Hd3a*) and *RICE FLOWERING LOCUS T 1* (*RFT1*), whose cognate proteins, commonly referred to as florigens, are essential to convert the vegetative SAM into a panicle (Komiya et al., 2008; Mineri et al., 2023). Florigens belong to the eucaryotic family of phosphatidyl ethanolamine-binding proteins (PEBPs) and have a general affinity to phospholipids (Nakamura et al., 2014; Serre et al., 1998). Upon expression in phloem companion cells, Hd3a and RFT1 are released into sieve elements through plasmodesmata, to reach distal plant tissues. Hd3a and RFT1 are loaded into the phloematic stream upon interaction with FT INTERACTING PROTEINS (FTIPs), including OsFTIP9 and OsFTIP1, respectively (Song et al., 2017a; Zhang et al., 2022a). Once in the SAM, the florigens interact with dimers of 14-3-3 proteins in the cytoplasm and migrate to the nucleus to form transcriptionally active complexes with DNA-binding proteins, including members of the bZIP family. These complexes promote the expression of genes required to form a panicle and are known as florigen activation complexes (FAC) (Cerise et al., 2021; Purwestri et al., 2009; Taoka et al., 2011; Tsuji et al., 2013). Thus, PPIs act at different levels during the floral transition and are coordinated with the environment in time and space.

At the SAM, Hd3a and RFT1 induce global changes of gene expression that eventually modify the overall plant architecture. Yet, the GRNs involved in such dramatic developmental switch have not been extensively explored (Gómez-Ariza et al., 2019; Mineri et al., 2023; Tamaki et al., 2015; Tsuji et al., 2015). Among the genes whose expression is induced at the SAM during the transition to flowering, *FLOWERING LOCUS T LIKE 1* (*FTL1*) is required to accelerate the conversion of the vegetative SAM into an IM, and to promote the transition from SBM to SM (Giaume et al., 2023). Mutations in *FTL1* delay flowering time additively when combined with *hd3a* or *rft1* mutants and increase the number of panicle secondary branches. Interestingly, *FTL1* transcripts and protein can be detected with an overlapping pattern, in all reproductive meristems from the IM to FMs, suggesting cell-autonomous functions.

Proximity labeling (PL) is a high-throughput technique that is being increasingly used for *de novo* discovery of PPIs and for reconstructing the proteome of subcellular compartments and cell types by means of organelle-/cell-specific protein baits (Kang & Rhee, 2022; H. B. Kim & Kim, 2024; Qin et al., 2021; Xu et al., 2021). This methodology was introduced over a decade ago but has been primarily applied to cell cultures and animal systems. During the last years, plant applications of PL have grown, including works on Arabidopsis, *N. benthamiana* and crop species, such as potato and tomato (Branon et al., 2018; Guo et al., 2023; Lee et al., 2024; Mair et al., 2019; Roux et al., 2012; Shi et al., 2023a; H. Tan et al., 2024; Tang et al., 2024; Xu et al., 2021).

PL is based on engineering a protein of interest (POI) by fusing it to a biotin ligase, resulting in biotin tagging of its interactors. Labelled proteins can be isolated from cell lysates using streptavidin-coated columns or magnetic beads. The resulting enriched biotinylated proteome is then analysed by mass spectrometry (MS). The advantages of PL include the ability of efficiently detecting unstable, transitory and compartment-specific PPIs, since protein tagging occurs *in vivo* and the integrity of protein complexes is not required for the protocol effectiveness. Biotinylation occurs on lysine residues that are present on any protein within 10 nm from the biotin-ligase fusion protein (or bait) (Kang & Rhee, 2022; D. I. Kim & Roux, 2016; Qin et al., 2021). Different versions of biotin ligases have been developed by directed evolution of the initial *E. coli* BirA enzyme, to further optimize biotinylation in eukaryotes. The TurboID (TbID) version differs from wild type BirA for 15 mutations, which permit promiscuous biotinylation of the POI interactome with faster kinetics and at room temperature (Branon et al., 2018; Roux et al., 2012).

In this study, we applied PL for the first time in rice stable transgenic plants, to study the proximal proteome of FTL1. We established a working PL protocol taking advantage of *N. benthamiana* as transient expression system. Subsequently, the technique was transferred to rice, where we globally analyzed biotin-tagged proteins in proximity to FTL1 by focusing on reproductive meristems at two different stages of panicle development. MS analyses on extracted proteins eventually returned lists of new potential interactors of FTL1, which suggest new properties during panicle development

## RESULTS

### Stability and biotinylation efficiency of FTL1-TbID in *Nicotiana benthamiana* leaves

To establish an efficient PL protocol to study the proximal proteome of FTL1, we conducted preliminary assays using a transient expression system. We selected the TurboID (TbID) as biotin ligase and used *N. benthamiana* leaves to test the expression, biochemical stability and biotinylation efficiency of FTL1-TbID fusions. To this aim, FTL1 was C-terminally fused to TbID, after its DNA sequence was optimized for the rice codon usage. C-terminal fusions were preferred over N-terminal, because several studies in different species have indicated that PEBPs retain their function when tagged C-terminally (Corbesier et al., 2007; Ho & Weigel, 2014; Kaneko-Suzuki et al., 2018; Komiya et al., 2008; Navarro et al., 2011; Taoka et al., 2011; Zhu et al., 2020). An HA tag was included after the TbID to allow immunological detection. Two variants of the fusion proteins were produced, in which the two moieties were separated by either a short or a long linking aminoacidic sequence (Fig. 1A). This was intended to examine whether a greater physical distance between FTL1 and TbID could affect protein function or labeling range. The spacing sequence was designed to acquire a disordered secondary structure and consists of 2xGGGGS in FTL1-*shortlinker*TbID (hereafter FTL1-*s*TbID) and 12xGGGGS in FTL1-*longlinker*TbID (hereafter FTL1-*l*TbID). A control construct containing the sole TbID (hereafter TbID), was also employed in these experiments. The promoter was adapted to suit the expression system and the species used. For expression in *N. benthamiana*, FTL1-TbID variants and TbID were placed under the control of the *CaMV 35S* promoter (Fig. 1A).

**Figure 1.**
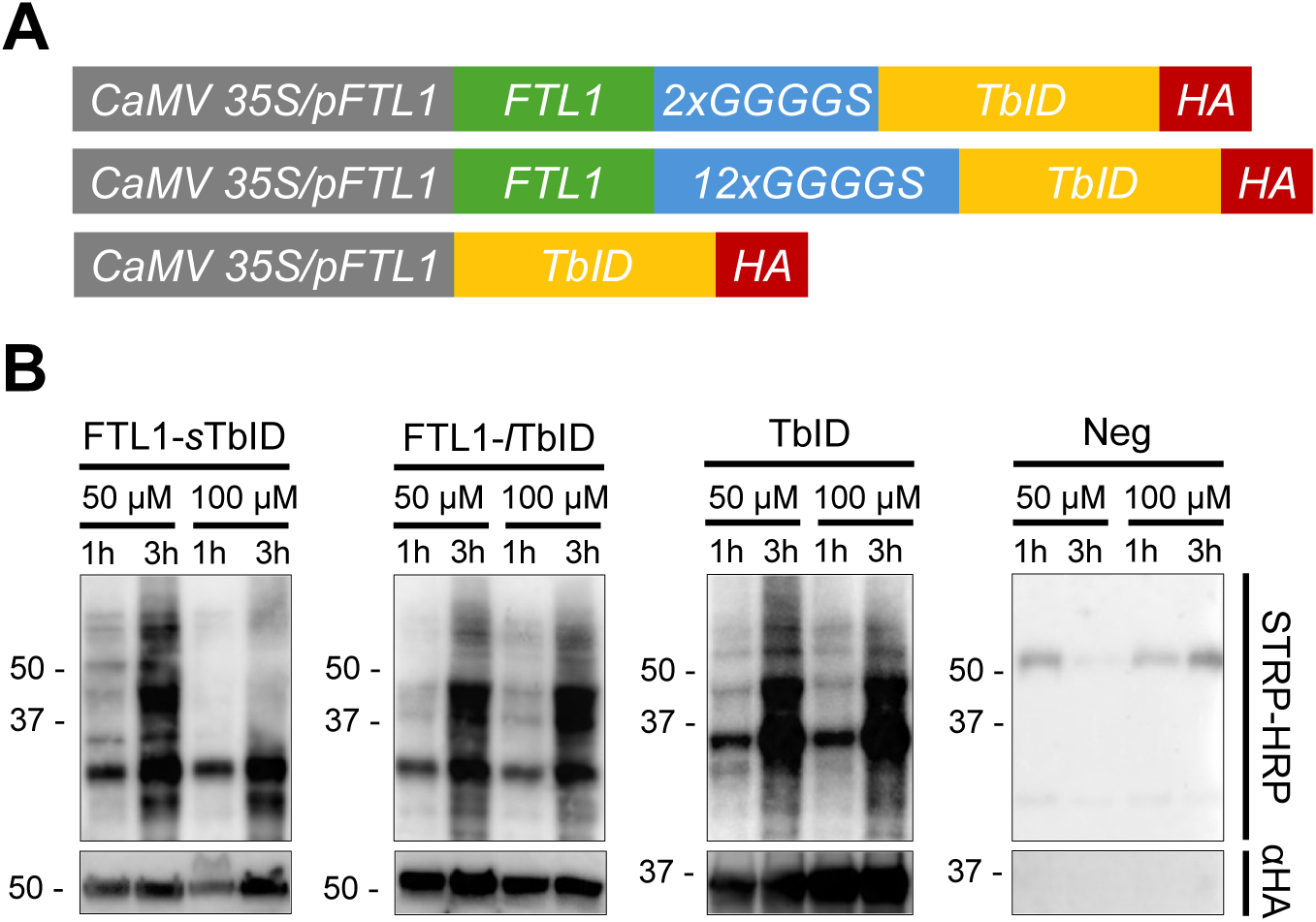
Expression and biotinylation efficiency of FTL1-TbID in *Nicotiana benthamiana*. Schematic representation of constructs used. FTL1 was C-terminally fused to TbID, separated by a short (2xGGGGS) or a long (12xGGGGS) linker formed by a repeated disordered sequence. An HA tag was included at the C-terminus of the enzymatic moiety. The TbID sequence was optimized for the rice codon usage and the start codon was removed from the TbID. In *N. benthamiana* assays, all proteins have been expressed under the CaMV 35S promoter sequence, whereas the pFTL1 promoter was used in rice. **B)** TbID-dependent biotinylation in agroinfiltrated *N. benthamiana* leaves. Agroinfiltrated leaves were treated with 50 μM or 100 μM biotin and sampled after either 1 or 3 hours. Western blots show the heterologous proteins (α-HA, anti-HA antibody, lower panels) and the biotinylated proteins (STRP-HRP, upper panels). The negative control (-) includes samples from a non-agroinfiltrated leaf treated with 50 μM or 100 μM biotin for 1 or 3 hours. Numbers on the left of the blots indicate protein molecular weight in kDa.

*N. benthamiana* leaves were individually agroinfiltrated with each construct, and western blots were performed on leaf protein extracts using an anti-HA antibody (α-HA), confirming the expression and stability of both the fusion proteins and the TbID (Fig. 1B). To observe the correlation between biotin availability over time and the amount of biotinylated proteins, exogenous biotin was provided to the agroinfiltrated leaves at two different concentrations (50 μM or 100 μM). Leaves were then sampled after 1 and 3 hours. Biotinylation activity was indirectly measured by western blot, using streptavidin conjugated to horseradish peroxidase (STRP-HRP, Fig. 1B). The biotinylation rate of transformed leaves was higher than in the non-agroinfiltrated control in each experimental condition. The pattern of biotinylated proteins varied according to sampling time, but it could not be correlated to the exogenous biotin concentration supplied. Indeed, a trend to higher labeling was observed in TbID and FTL1-TbID sampled 3 hours after exogenous biotin treatment, both at 50 and 100 μM concentration. No substantial differences were observed between leaves that were treated with different biotin concentrations but sampled at the same timepoint. This suggests that the activity of the TbID enzyme was maximum even at the lowest biotin concentration, and that time is the major parameter positively correlated with biotinylation in *N. benthamiana* leaves. Overall, these preliminary tests indicate that both the two FTL1-TbID versions and TbID are stably expressed in agroinfiltrated *N. benthamiana* leaves and retain high enzymatic activity.

### Streptavidin-mediated precipitation allows effective enrichment of biotinylated proteins

To complete the working PL pipeline in *N. benthamiana*, we performed purification of biotinylated proteins followed by mass spectrometry (MS). To this end, *N. benthamiana* leaves were agroinfiltrated with TbID and the two FTL1-TbID construct variants. Biotin was provided at 50 µM concentration for 1 hour, to reduce overlabeling. Leaf protein lysates were first filtered through ZebaSpin columns, to eliminate free biotin molecules that could negatively affect the subsequent enrichment steps. Next, the labelled proteome was pulled down from the total extract using streptavidin-coated beads and the resulting protein precipitates were checked with silver staining (Fig. 2A). Protein bands were visible in all agroinfiltrated samples and in the negative control, consisting of a leaf to which only biotin was provided. After pull down, the TbID precipitate showed more bands compared to the two FTL1-TbID versions and the negative control, indicating extensive non-specific labeling (Fig. 2B). Western blot on the precipitated proteins, using STRP-HRP, confirmed that the TbID samples were the most enriched in biotinylated proteins, while FTL1-*s*TbID and FTL1-*l*TbID pull downs presented a high number of bands that were not visible in the negative control. Almost complete depletion of biotin-labelled proteins in the flow-throughs confirmed optimal binding by streptavidin-coated beads (Fig. 2B).

**Figure 2.**
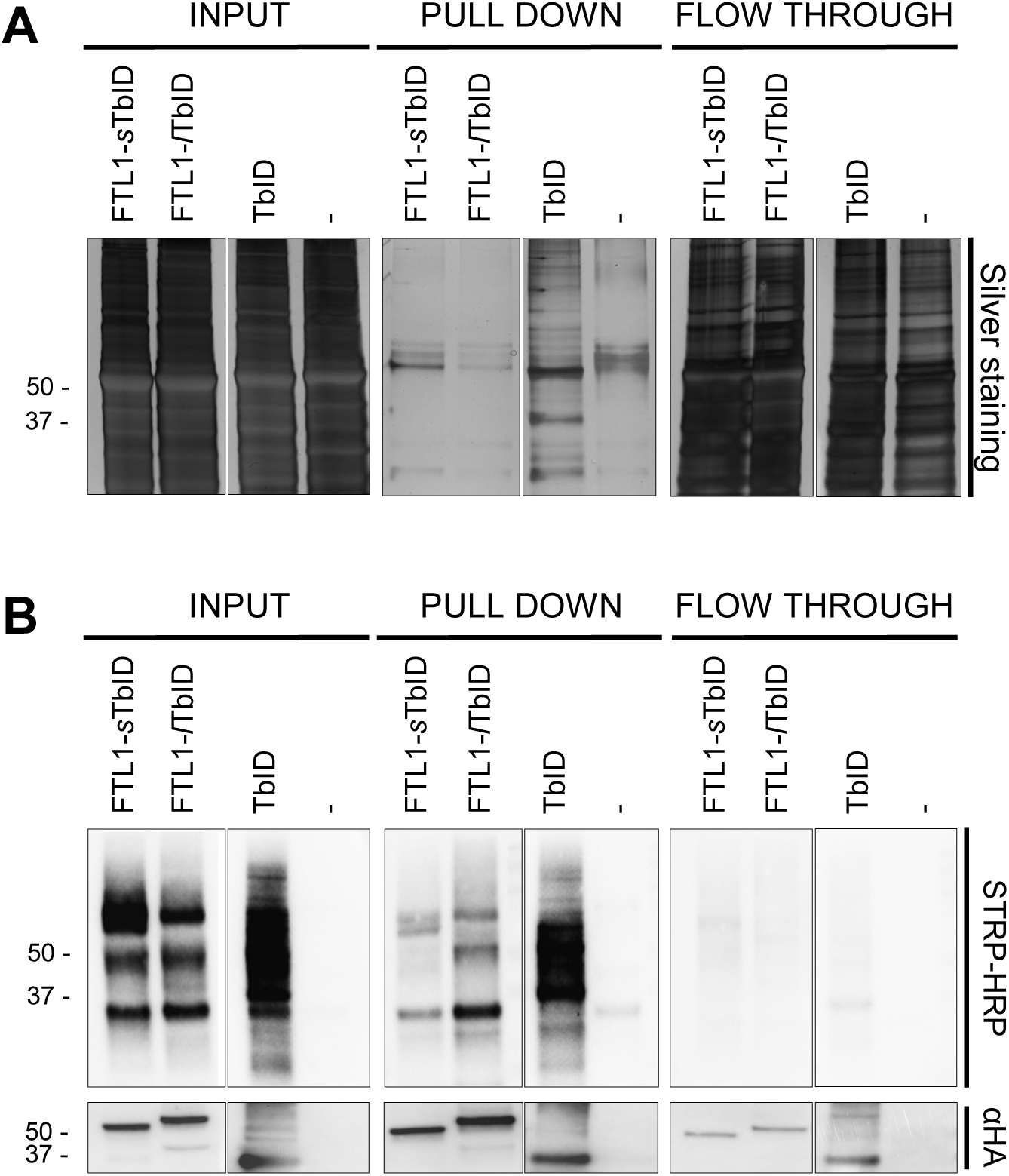
Precipitation of *N. benthamiana* biotinylated proteins. **A)** Silver staining of protein extracts from *N. benthamiana* leaves, sampled 1 hour after 50 µM biotin infiltration. The left panel shows the total proteome (input), whereas the middle and right panels show the pull downs and flow through, respectively. Pull downs were enriched for biotinylated proteins using STRP-coated beads. **B)** Western blots of the samples in **A)** using STRP-HRP (upper panels) or anti-HA (lower panels) antibody. The number of biotinylated proteins is highest in the TbID and minimum in non-infiltrated leaves.

Next, pull downs of agroinfiltrated *N. benthamiana* leaves were analyzed through MS to obtain proteomic datasets about non-specific biotinylation targets of TbID and the effect of linker length on the labeling rate of FTL1-TbID (Supplementary Table 1). This experiment was primarily aimed at assessing features of the FTL1-TbID fusions, as well as unraveling any technical challenge. However, as FTL1 is a close homolog of NtFT4 (78% identity), the main tobacco florigen (Harig et al 2012), we reasoned that we could also gather biologically relevant candidates from the proximal proteome of FTL1 in tobacco. Precipitates were analyzed in triplicates and the peptides were searched over the reference proteome of *N. benthamiana*, as well as that of its close relative *N. tabacum*. The latter is better annotated, ensuring a higher number of identified peptides per sample, and is accessible to GO Enrichment Analysis by the Gene Ontology consortium.

In our analyses, a protein was considered enriched compared to the control if it had an absolute fold change (FC) > 2 and a false discovery rate (fdr) < 0.05 (enriched hit), or if it had an absolute FC > 1.5 and fdr < 0.02 (enriched candidate). In TbID, 29 *N. benthamiana* and 225 *N. tabacum* proteins were found as enriched against the negative control, consisting of the pull downs of non-agroinfiltrated leaves, after biotin injection (Supplementary Table 1). According to the *N. tabacum* ontological analysis, more than 40% of identified proteins had basal metabolic functions, belonging to primary and phenylpropanoid/isoprenoid pathways. The assigned categories of translation, protein folding and protein degradation accounted together for another 15% of the total enriched proteome (Supplementary Table 2, Fig. 3A). We believe that these categories include non-specific interactors due to biotinylation of proximal proteins during TbID translation and processing. Such a functional distribution is also mirrored by protein localization, with almost 40% of the enriched proteins being exclusively cytosolic, and 14% being present in two or more cell compartments (ubiquitous). Less than 4% of proteins in the list were nuclear (Supplementary Table 2, Fig. 3B).

**Figure 3.**
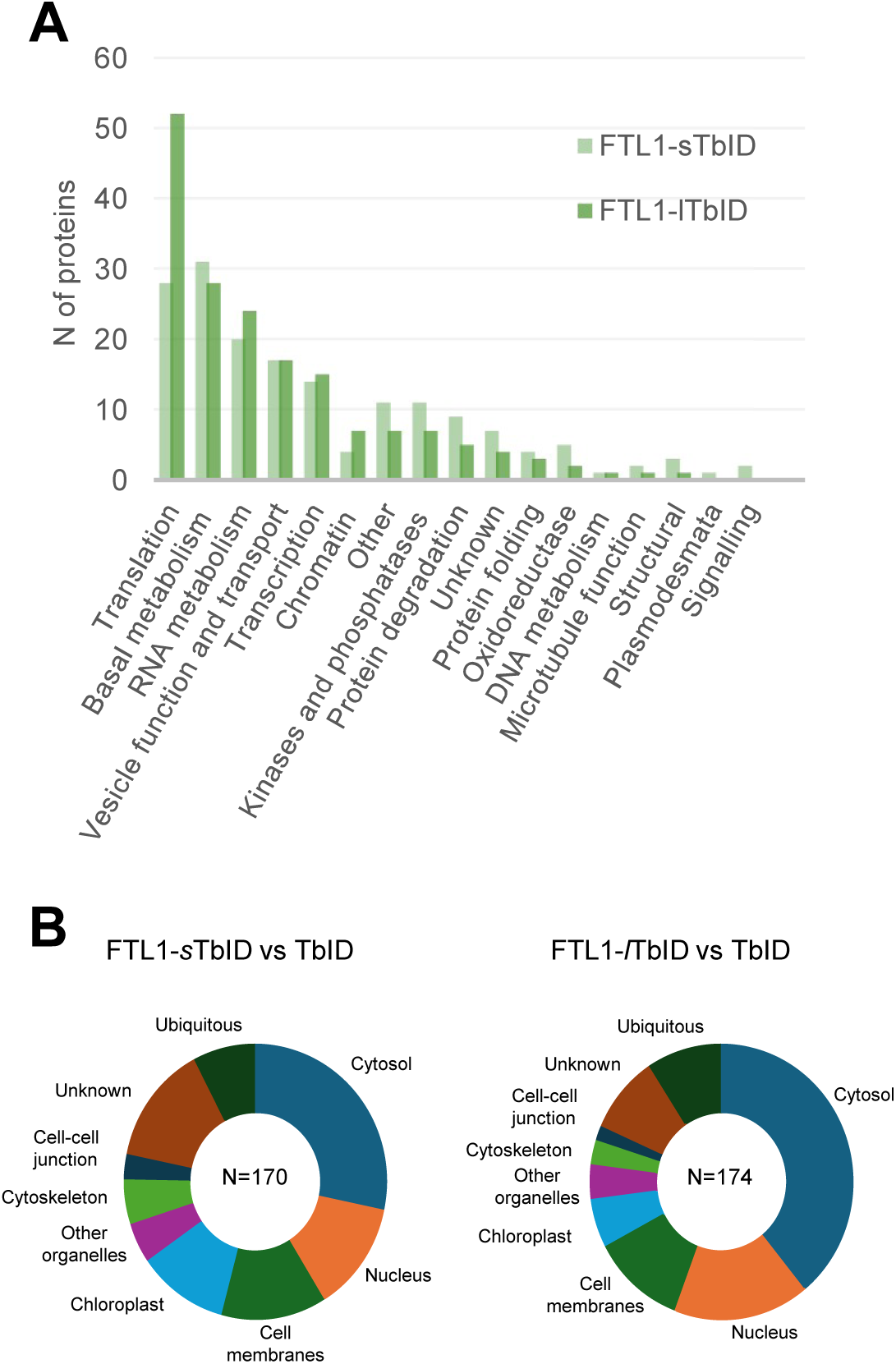
The proximal proteome of FTL1 in *Nicotiana.* **A)** Graphic bar chart representation of the FTL1 interactome in *N. tabacum*. The enriched elements of FTL1-*s*TbID and FTL1-*l*TbID PL datasets are grouped according to their predictive biological function. **B)** Pie chart representation of the predicted subcellular localizations of the proteins represented in A).

Compared to TbID, 17 and 22 *N. benthamiana* proteins were enriched in FTL1-*s*TbID and FTL1-*l*TbID, respectively, being 10 of them common to both (Supplementary Table 1). Since the number of enriched proteins in FTL1-*s*TbID was only slightly lower than in FTL1-*l*TbID, it cannot be sustained that the linker length poses substantial differences in the amount of biotinylation events taking place in the bait surroundings. However, the non-complete overlap of the two datasets indicates that the nature of the PPIs may possibly change depending on linker length.

The protein interactors common to FTL1-*s*TbID and FTL1-*l*TbID in *N. benthamiana* include the autophagy cargo receptor NEIGHBOUR OF BRCA1 (NBR1) and a germin-like protein with a predicted localization to plasmodesmata (Table 1). The degree of enrichment of the proteins varied between the two bait versions, and in general, no pattern of enrichment was evident that could suggest distinct properties of long versus short linker.

**Table 1.**
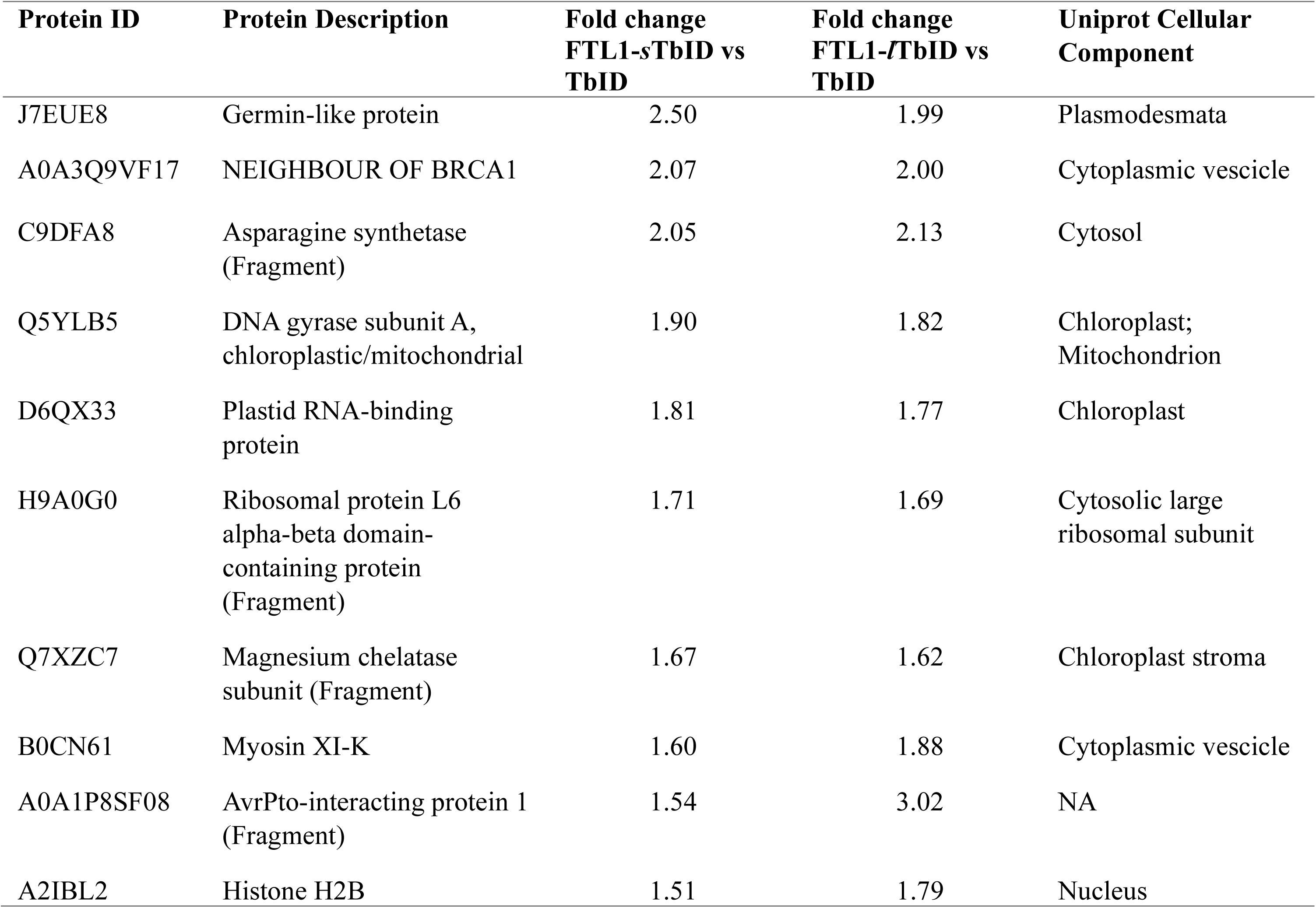
List of FTL1 proximal proteins common to FTL1-*s*TbID and FTL1-*l*TbID in *N. benthamiana*.

The research over the *N. tabacum* reference proteome delivered 183 and 185 potential interactors for FTL1-*s*TbID and FTL1-*l*TbID, respectively, being 106 in common to both (Supplementary Table 2). Among those, more than 50% are involved in transcription, RNA metabolism, basal metabolism or vesicle function and transport. Another abundant category is related to translation, even if remarkably more enriched in FTL1-*l*TbID (29%) than in FTL1-*s*TbID (16%), which could be partially explained by an increase of the labeling range of FTL1-*l*TbID during its translation (Fig. 3A). The FTL1-*l*TbID interactome displayed a higher proportion of cytosolic and ubiquitous proteins if compared to FTL1-*s*TbID. Conversely, the latter was richer in proteins localized to the cytoskeleton, chloroplasts and the intercellular junction (including plasmodesmata and the extracellular matrix). In the two datasets, an equal fraction of proteins is assigned to the nucleus and to cell membranes, each exceeding 10% of the total interactome (Fig. 3B). Overall, these results indicate that the PPI pattern of FTL1-TbID is substantially distinct from that of TbID and might depend on conserved biochemical properties governing FTL1 affinity to given protein types and specific plant cell structures.

### Expression of FTL1-TbID in rice causes early flowering and increases total protein biotinylation in developing panicles

To study the proximal proteome of FTL1 during rice flowering, FTL1-*l*TbID and FTL1-*s*TbID were stably transformed into rice plants under the control of the *pFTL1* endogenous promoter, by *in vitro* transformation of *O. sativa* var. Nipponbare calli. Primary FTL1-*l*TbID transformants exhibited extreme phenotypes: the plants were extremely reduced in size, flowered early (within one month from transfer to soil) and generated small sterile panicles with elongated bracts harbouring sterile flowers, ultimately resulting in no seeds (Fig. 4A-C). In contrast, FTL1-*s*TbID transgenic lines retained normal plant size and fertility (Fig. 4B). We selected two independent FTL1-*s*TbID transgenic lines (#1.3 and #2.5) and measured their flowering time. Both lines were early flowering both under SD and LD conditions. However, they flowered later under LD compared to SD, indicating that photoperiod sensitivity was retained (Fig. 4A).

**Figure 4.**
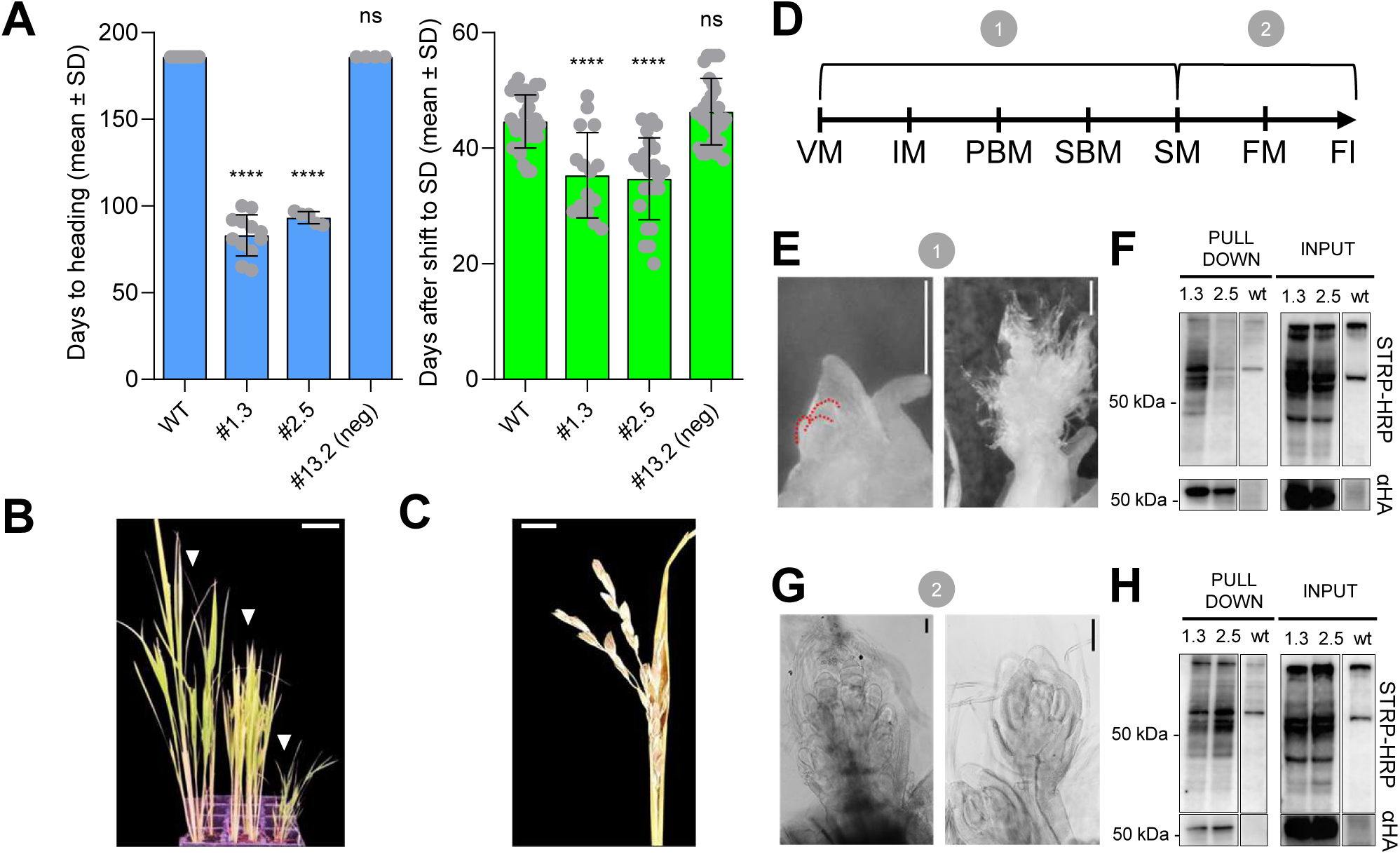
FTL-TbID transgenic rice lines flower earlier and show increased protein biotinylation in flowering meristems. **A)** Flowering time of *pFTL1:FTL1-sTbID* plants grown under LD (left) or under LD for four weeks and then shifted to SD (right). Flowering time is expressed in days from sowing to heading for the LD experiment, and as days from shifting to SD to heading for the SD experiment. Line #13.2 (neg) is a transformation escape not harboring the transgene. The LD experiment was interrupted after 180 days, before flowering of the negative control lines. Each dot represents a plant. Asterisks indicate statistical significance based on two-tailed Student’s *t* test (****, p<0.0001; ns, non-significant). The experiment was repeated three times independently, with similar results. **B)** Phenotype of *pFTL1:FTL1-TbID* transgenic rice plants at flowering. From left to right: wt, *pFTL1:FTL1-sTbID* and *pFTL1:FTL1-lTbID* plants. **C)** Sterile panicle of a representative *pFTL1:FTL1-lTbID* line. **D)** Schematic representation of the progression of panicle development from the vegetative meristem to flowers. One and two indicate the sampled stages. **E)** Representative pictures of stage 1 samples, including panicles at PBM (left) and SM (right) stages. **F)** Western blots of stage 1 samples from inputs and pull downs using STRP-HRP (upper panels) or α-HA antibody (lower panels). **G)** Representative pictures of stage 2 samples, including panicles at FM (left) and flower (right) stages. **H)** Western blots of stage 2 samples from inputs and pull downs, using STRP-HRP (upper panels) or α-HA antibody (lower panels). Scale bars in B), 10 cm, in C), 1 cm, in E) and G), 200 µm.

Next, we used lines #1.3 and #2.5 to test FTL1-TbID stability and its biotinylation efficiency. To this extent, plants were grown for 1 month under LD and induced to flower upon shifting to SD conditions. We harvested developing panicles at two stages of differentiation, following selection at the stereomicroscope. In this study, stage 1 corresponds to early panicle development and includes meristems ranging from IM to SM (Fig. 4D, E). Stage 2 corresponds to late panicle development and includes meristems ranging from SM to FM (Fig. 4D, G). To reduce excessive biotinylation and unspecific labeling, no exogenous biotin was given to the plant tissues before sampling. Expression of FTL1-*s*TbID in stage 1 and 2 panicles was confirmed by western blot (Fig. 4F, H), and the IP on the total biotinylated proteins was performed as described for *N. benthamiana*. Despite adding no extra biotin, the cell lysates of both panicle stages contained more biotinylated proteins than the wt, providing indirect evidence of FTL1-*s*TbID enzymatic activity (Fig. 4F, H). Silver staining on STRP-precipiated protein extracts also showed visible, specific bands in stage 1 and stage 2 pull downs (Supplementary Fig. 2). These experiments indicate that stage 1 and 2 panicles of FTL1-*s*TbID, lines #1.3 and #2.5, are suitable for protein analysis and had the prerequisites for PL implementation in rice.

### PL defines the proximal proteome of FTL1 at two distinct panicle differentiation stages

Proteomic analyses were carried out on the biotinylated proteins pulled down from stage 1 and stage 2 panicles of FTL1-*s*TbID #1.3 and #2.5 lines, using the wt as a control. The same pipeline used for *N. benthamiana* PL experiments, including sample processing, MS and data analysis, was also applied here.

PL implementation produced two datasets of more than 3000 proteins each (Supplementary Table 3). In stage 1, 72 proteins were enriched against the control in at least one of the two transgenic lines, whereas 26 were common to both lines. Stage 2 dataset was characterized by a higher protein enrichment, with 197 potential FTL1 interactors identified in at least one of the two lines, and 102 common to both lines. Only proteins that resulted enriched in both independent transgenic lines in each stage were considered in subsequent analyses (Supplementary Table 4). In every dataset, FTL1-*s*TbID was also found enriched, as expected from its self-biotinylation.

The biological function categories assigned to the enriched proteins revealed a diverse distribution between the two stages, mainly due to the great difference in protein abundance, with many categories resulting under-or non-represented in stage 1. For example, RNA metabolism and chromatin categories were almost exclusive of stage 2, representing together more than 30% of the enriched proteins for this stage. Also, categories associated to protein synthesis and processing were found only in this stage (Fig. 5A).

**Figure 5.**
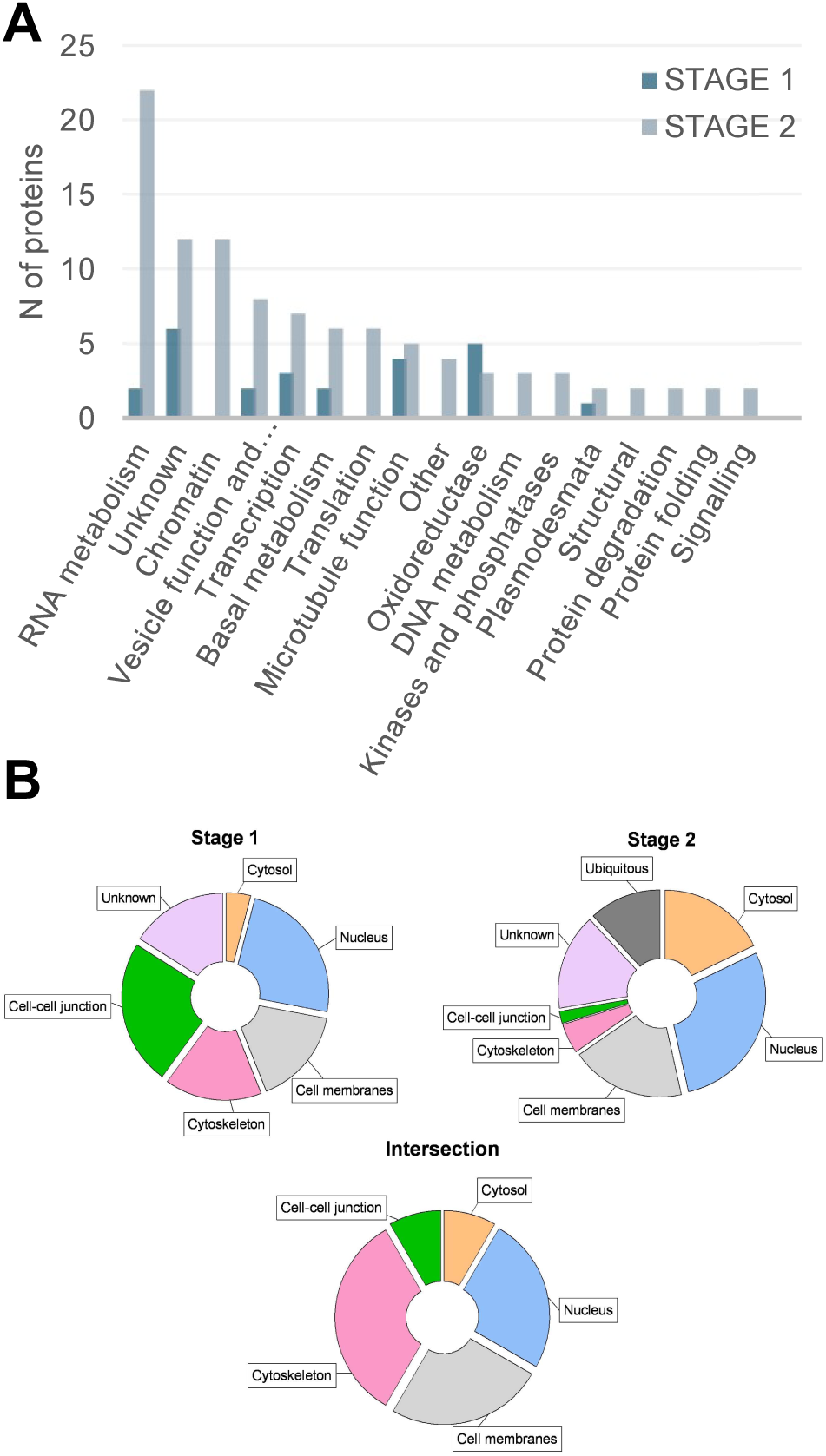
The proximal proteome of FTL1. **A)** Biological functions of FTL1 interactors. The bars indicate the number of interactors identified in stage 1 and 2, based on the functional categories defined in PANTHER. When not available, categories were manually curated, upon protein database searches. **B)** Cellular localization of FTL1 interactors. The pie charts represent the compartments assigned to FTL1 interactors identified at stage 1 (left), stage 2 (right), or common to both stages (bottom).

In terms of cell localization, more than 50% of the enriched elements were found in the nucleus, in cell membranes or in the intercellular junction in both stages. Proteins assigned to the cytoskeleton, comprising microtubular elements, were proportionally more abundant in stage 1. These four categories were predominant in the intersection between stage 1 and 2, making more than 90% of the list (Fig. 5B).

Although stage-specific interactors may be present in the lists, the overlap between stage 1 and 2 identified 12 proteins that stand as the most probable FTL1 interactors during panicle differentiation (Table 2). These include the transcription factors Auxin Response Factor 12 (ARF12) and NPR-1 like 5/BLADE-ON-PETIOLE 1 (OsNPR5/BOP1), four microtubule-associated proteins, one of which predictively localized to the plasmodesmata (LOC_Os06g17440), and three components of the vesicular trafficking pathway.

**Table 2.**
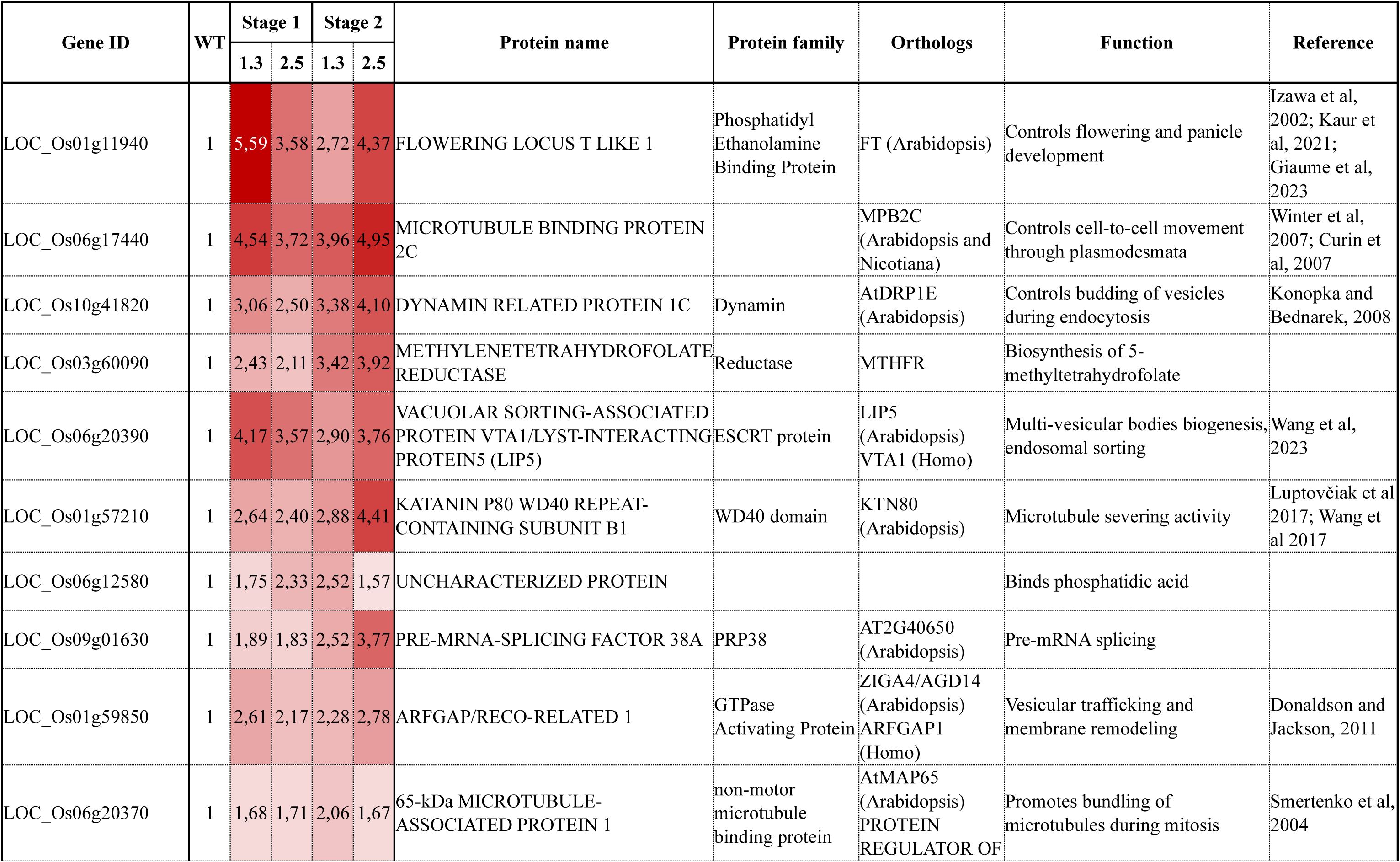
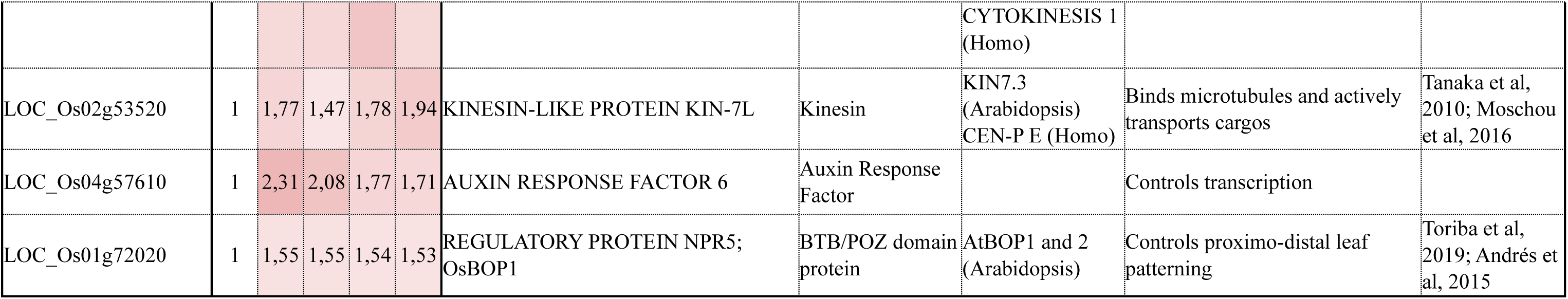
List of FTL1-*s*TbID proximal proteins common to stage 1 and stage 2 panicles in *O. sativa*.

Among the stage-specific potential FTL1 interactors, MAINTENANCE OF MERISTEM LIKE 1 (MAIL1, stage 1), HEAT SHOCK FACTOR A1 (HSFA1), OsSWEET1A and OsFCA (stage 2) can also be encountered. Homologues of these proteins have been reported to be involved in thermomorphogenesis, flowering and tuberization in different species (Abelenda et al., 2019; Andrés et al., 2020; Lauschke et al., 2025; Macknight et al., 1997; Mineri et al., 2025; Mishra et al., 2002; W. Tan et al., 2025; Toribio et al., 2024; Yan et al., 2010). OsGLP5-2, a germin-like protein putatively localized to plasmodesmata, is also enriched at stage 2, and is orthologous to the germin-like protein identified in the *N. benthamiana* PL experiment. This seems to hint at a conserved affinity of FTL1 to transmembrane germin-like receptors. Interestingly, we identified five CCCH zinc finger proteins specific to stage 2, none of which has been characterized in rice. The CCCH class includes proteins that can bind either DNA or RNA and is involved in several biological processes.

### FTL1 protein shows cell-to-cell mobility properties like rice florigens

The identification of several proteins associated to microtubules or vesicular trafficking at both stage 1 and 2, of a germin-like protein in both stage 2 and *N. benthamiana*, and of a microtubule binding protein involved in cell-to-cell trafficking through plasmodesmata, suggested that FTL1 could be possibly transported from cell to cell through plasmodesmata. The expression patterns of *FTL1* transcripts and protein suggest cell-autonomous function, but local movement between meristematic cells of the developing inflorescence cannot be excluded. To test FTL1 protein mobility, we performed assays in *N. benthamiana*, as described in Ohtsu et al. (2024) To this end, the *FTL1* nucleotide sequence was fused to *GFP* at the C-terminus and placed under the control of the *CaMV 35S* promoter. The same vector contains a nuclear localized dTomato, also expressed by the *CaMV 35S* promoter. Agroinfiltration was carried out at a very low cell density (OD600 of 3×10^-5^) to identify isolated, individually transformed cells, as visible by a single red fluorescent nucleus (Fig. 6A). The mobility degree of the target protein was measured as the number of surrounding cells that presented a green signal, but not a nuclear-localized red one. To get a comparison with FTL1 homologues known to move over long distances, Hd3a and RFT1 mobility was also tested with the same approach. GFP and 2xGFP were used as positive and negative controls, respectively, since a single, but not a double, GFP can move through plasmodesmata (Ohtsu et al., 2024).

**Figure 6.**
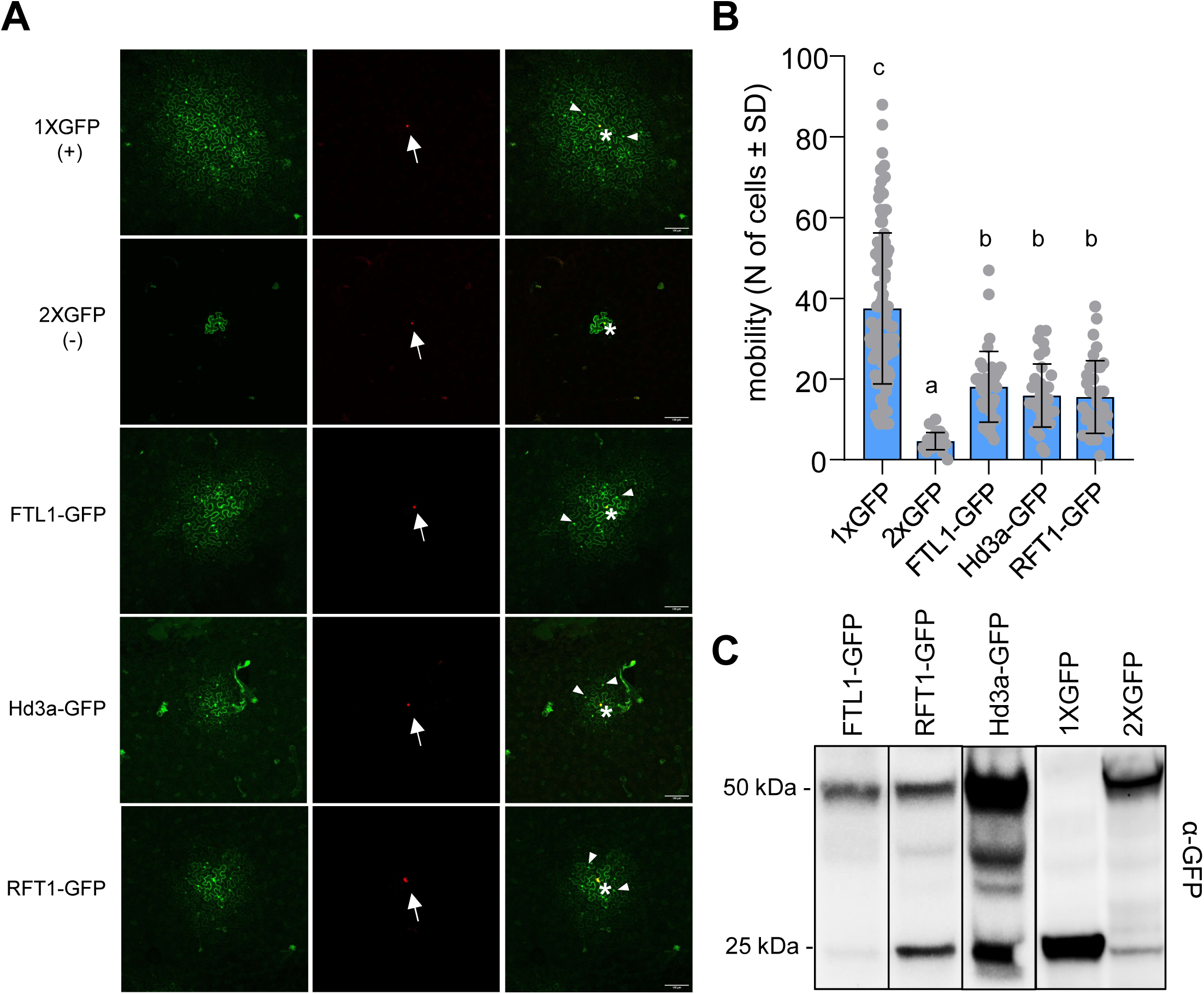
Mobility assays of FTL1, Hd3a and RFT1 in *N. benthamiana* leaves. **A)** Confocal imaging of representative cells used in the assays. The red signal indicates dTomato fluorescence, whereas green indicates GFP. Isolated cell-transformation events are highlighted by a white arrow pointing to a single fluorescent nucleus in the NLS-dTomato pictures, and by a white star in the merged. White arrowheads represent cells the GFP has moved into. **B)** Graphical representation of the mobility assays. The intercellular mobility of single GFP (mobile control), 2xGFP (non-mobile control) and GFP-tagged PEBP proteins is measured as the number of cells expressing GFP but not NLS-dTomato in the surroundings of a single NLS-dTomato-expressing cell. Each dot represents an independent cell expressing dTomato and GFP/protein-GFP. Statistical analysis has been performed using a Kruskal-Wallis nonparametric test, with a Dunn test as a post-hoc test. **C)** Western blots performed on infiltrated tobacco leaves using anti-GFP antibody.

This assay showed that FTL1 was able to move to non-infiltrated cells similarly to the main rice florigens Hd3a and RFT1 (Fig. 6A, B). Western blots were performed on leaf total protein extracts to confirm that the detachment of the fluorophore moiety did not take place more frequently than in the negative control, unwantedly generating mobility artefacts (Fig. 6C). The non-mobile 2xGFP and the PEBP-GFP proteins have a similar molecular weight (∼50 and ∼45 kDa, respectively), but while 2xGFP is retained in the infiltrated cell, PEBP-GFP move to adjacent cells, suggesting that their transport is not passive but active, and mediated by factors that can modulate the size exclusion limit of the plasmodesmata.

## DISCUSSION

Plant developmental processes are highly plastic and dynamic and often involve fast changes in PPIs to take place correctly. In recent years, proximity labeling has been increasingly used in plants to identify the interacting partners of a protein of interest. While most studies have been performed in Arabidopsis stable transgenic lines, examples of PL applications in other species, including tomato and tobacco, are available (Arora et al., 2020; Mair et al., 2019) The predominant use of dicot species is justified by the ease of transformation (often transient), fast cycling and non-limiting sampling material, which allows testing multiple conditions simultaneously. Instead, PL in rice has been reported by Lin et al. (2017), only using transient expression in protoplasts and exploiting a first-generation biotin ligase, whose efficiency is lower compared to newly available third-generation enzymes (Q. Lin et al., 2017). Taking full advantage of PL requires expanding the range of species towards monocots, adapting the system while considering technical aspects already explored in dicots.

In this research, a working methodology for PL application in rice has been developed, using the FTL1 florigen-like protein as bait. FTL1-TbID was introduced into *N. benthamiana* leaves transiently through agroinfiltration, while being stably transformed into rice transgenic lines to gain novel insights into FTL1 function in developing panicles.

Our data indicate that FTL1 is proximal to several proteins, including components of the cytoskeleton, vesicular trafficking proteins, as well as regulatory proteins. While some of the identified interactors might be more generally involved in FTL1-TbID translation, processing and intracellular transport, others might be, instead, more directly linked to FTL1 function and mobility to adjacent cells during the SAM reproductive differentiation.

### Establishing a proximity labeling protocol in rice

Preliminary experiments in *N. benthamiana* leaves were instrumental to determine the biochemical parameters to be used for the application of PL on FTL1 in rice. The use of complementary protein analysis techniques, including western blot and silver staining, were necessary to get an indirect description of the samples before employing “omics” methods.

The first issues to be considered in every PL experiment include the choice of the biotin ligase and the exogenous supply of biotin. In plant cells, biotin is naturally present in concentrations usually below 13 μM, with a slight variability depending on cell type and compartment (Alban, 2011; Alban et al., 2000; Baldet et al., 1993). The affinity to biotin of the BioID ligase, a precursor of TbID, has been reckoned to a Km = 0.3 μM, suggesting that endogenous biotin would suffice for protein tagging in plants (Arora et al., 2020). However, in published studies, biotin was provided at concentrations ranging from 50 μM to 250 μM. Under saturating biotin concentrations, time is the factor that correlates most to promiscuous biotinylation, and the principal parameter to consider for the control of the general labeling extent in PL experiments (Arora et al., 2020; Branon et al., 2018; P. Li et al., 2017; Mair et al., 2019; Roux et al., 2012). Biotinylation of surrounding surface-exposed lysines by TbID occurs within a range of 10 nm, in less than 1 h (T.-W. Kim et al., 2023; Mair et al., 2019; Samavarchi-Tehrani et al., 2020). Longer exposure to externally supplied biotin may be required for low abundant proteins, such as transcription factors, but it can be counterproductive since false positives could increase dramatically and free biotin could interfere with further sample processing (Arora et al., 2020; T.-W. Kim et al., 2023; Mair et al., 2019; Xiong et al., 2021; Xu et al., 2021). In our tobacco experiments, we used high biotin concentrations which were meant to balance overexpression of the biotin ligase driven by the *35S* promoter. However, in rice, we decided not to treat samples with extra biotin. The main reason behind this choice was to limit the number of false positives. The faster kinetics of TbID, compared to BioID or miniTbID, suggests that TbID could be particularly suitable for effective interactor labeling in the absence of external biotin (Branon et al., 2018; Guo et al., 2023; May et al., 2020; Xiong et al., 2021). Additionally, the labeling efficiency of TbID was shown to be similarly high both at 22°C and 30°C, the latter corresponding to rice growth temperature (Mair et al., 2019). Finally, since our starting material included rice inflorescence meristems at two developing stages (differently from previous studies that mostly employed whole Arabidopsis seedlings), avoiding biotin treatments allowed faster sample processing and tissues integrity. Our preliminary tests on *pFTL1:FTL1-sTbID* rice lines confirmed these premises, avoiding unwanted environmental stress. However, it cannot be excluded that some low abundance proteins normally associated to FTL1 might not have been detected in our analyses due to higher biotin needs.

An appropriate PL bait should behave as much as possible like the POI in its natural context. Therefore, vector construction should be adapted to the specific protein being studied. The choice of N-versus C-terminal fusions, linker length, additional tags and the promoter driving expression of the POI, can influence the outcomes. The early flowering phenotype shown by our transgenic lines indicated that the main function of FTL1 was preserved when TbID was C-terminally fused to it, despite the fusion protein having almost a double size than the wild type. Early flowering might depend on a dose-dependent effect of the bait in a wild-type background already encoding a functional FTL1. Alternatively, the *pFTL1* endogenous promoter driving expression of FTL1-TbID might be lacking some repressive elements. Nevertheless, photoperiod sensitivity was retained, at least in *pFTL1:FTL1-sTbID* which was an essential requisite in our experimental system. Complementation of the mutant by the POI is the best strategy to assess functionality of the TbID-tagged protein (Mair et al., 2019). However, in the case of *FTL1*, available mutants were produced by CRISPR, and still express Cas9 and gRNAs that would target the *FTL1-TbID* locus (Giaume et al., 2023). Extremely early flowering and aberrant reproductive development in *pFTL1:FTL1-lTbID* lines may be explained by exacerbation of protein function linked to changed molecular features in presence of a longer disordered linker, although there is no additional data backing this hypothesis. However, we could compare the performance of two different linker lengths in tobacco. Based on our data, having a longer spacing sequence is not directly related to higher biotinylation rates, but rather contributes to changing the interaction properties of the bait. In the long-linker FTL1-TbID, translation-related, RNA metabolism, ribonucleoproteins and splicing factors targets increased, and their biotinylation may simply be due to bigger bait size. These functional categories could easily encompass non-specific TbID targets and may represent false positives also in the FTL1-sTbID vs TbID comparison, since the fusion protein is bigger in size than the control. Similar categories were also enriched in other studies (X. Li et al., 2023; Mair et al., 2019), and may represent common false positives to be expected in plant datasets. Nonetheless, it is remarkable that a small percentage of the enriched proteins in *N. benthamiana* is common to both FTL1-TbID versions, raising the issue of their biological value. Finally, the insertion of an HA tag, rather than a bigger GFP or YFP tag, as reported in other studies, contributed to maintain the size of FTL1-TbID as close as possible to endogenous FTL1 (H. B. Kim & Kim, 2024; X. Li et al., 2023; Mair et al., 2019; Wallner et al., 2024). While a fluorescent tag might be helpful to assess POI tissue specificity, its size might, in fact, interfere with the protein function.

Once the molecular prerequisites for PL on FTL1 were met, we implemented an extraction and protein precipitation protocol that could fit both *N. benthamiana* and rice. The use of a harsh denaturing buffer (see Methods) is fundamental to completely lyse all subcellular compartments and ensure spatial sensitivity of the proteomic analyses (Guo et al., 2023; Xu et al., 2021). In our experiments, binding of labelled proteins by streptavidin-coated beads was not impaired by the extraction buffer composition that can be challenging when working with antibodies. In fact, the high strength of the streptavidin-biotin bond, which has been calculated as K_d_=10^-14^ mol/L, makes the affinity purification protocol extremely efficient (Fridy et al., 2015; Hou et al., 2016). Additionally, the great number of precipitated proteins delivered in our pull downs indicate that beads washing and protein elution steps were also performed under optimal conditions. Possible protein losses during desalting, executed prior to overnight pull-downs, have not been calculated. This step is fundamental to maximize protein binding to streptavidin beads, especially in tissues containing elevated amounts of biotin, which could saturate the beads and reduce their affinity to the preys, or when the number of biotinylated proteins is expected to be limited. To this purpose, the use of ZebaSpin columns, instead of the widely employed PD10 filters, better fit our working volumes reducing waste risks (Arora et al., 2020; Grismer et al., 2024; Z. Li et al., 2022; Z. Lin et al., 2024; Shi et al., 2023b).

Experimental design can also contribute to reducing the number of false positives. In rice, the use of a wild type control allowed the removal of beads contaminants or naturally biotinylated proteins, but not those that may be common targets of the TbID. For example, acetyl-CoA carboxylases and methylcrotonyl-CoA carboxylases, which are biotinylated enzymes, were highly abundant in wt samples, as previously reported (Q. Lin et al., 2017). If compared to *N. benthamiana/N. tabacum* datasets, the percentages of candidate FTL1 interactors vs the total detected proteome in rice meristems analyses is very low, suggesting high labeling specificity. Finally, the use of two independent transgenic lines and two distinct developmental stages further contributed to restricting the list of candidate interactors to the most probable ones.

At a proteomic level, the most favorable sensitivity was conveyed using multiplexed Tandem Mass Tag (TMT) mass spectrometry, where multiple samples are labelled with unique tags and analyzed simultaneously, increasing throughput and reducing technical variability.(Arul & Robinson, 2018; Pappireddi et al., 2019; Rakk et al., 2023). A statistical analysis implying two contiguous thresholds of differential abundance and significance helped in identifying the candidate interactors (see Methods). By keeping a high significance threshold for abundant proteins (FC > 2, enriched hits) potential false positives were excluded, whereas less abundant interactors (FC > 1.5, enriched candidates) were identified by decreasing the lower significance limit.

The distinct degree of protein enrichment between stage 1 and stage 2 datasets can be due to differences in the total proteome of the two developmental stages, or to specific changes of FTL1 function. Indeed, some candidate interactors can be stage specific. However, a reduced number of enriched proteins for stage 1 could be very likely due to the greater technical variance for this experiment, since each sample comprised twice the meristems than stage 2, and could have occasionally contained off-target contaminants from close plant tissues, given that they were smaller and more difficult to sample.

### FTL1 proximal interactors suggest its involvement in distinct cellular pathways

Our proximal proteome analysis did not list any known component of the flowering activation complex (FAC) at any stage of panicle development tested. OsFD and 14-3-3 proteins are expressed in the SAM, and their interaction with PEBP proteins, including FTL1, has been demonstrated by several research groups (Cerise et al., 2021; Ding et al., 2016; Giaume et al., 2023; Jang et al., 2017; Kaneko-Suzuki et al., 2018; Kaur et al., 2021; Purwestri et al., 2009; Taoka et al., 2011; Tsuji et al., 2013). Inaccessibility of lysines and the presence of the TbID tag may have impeded successful biotinylation of FAC components. A previous PL experiment in transfected rice protoplasts using BirA, the first available biotin ligase, identified multiple 14-3-3 proteins as potential interactors of OsFD1/2, but did not identify PEBP proteins, likely because of their absence in seedlings-derived protoplasts (Q. Lin et al., 2017). Alternatively, although PL on OsFD1/2-BirA involved constitutive overexpression of the baits and 24-hours exposure to saturating biotin concentrations, low-abundant proteins such as the PEBPs might be below the detection limits of the method. A similar problem could have occurred with FTL1-TurboID in developing panicles, particularly if the relevant PPIs take place in a limited number of cells and for a short time. However, FTL1-TbID conserved its main flowering promoting function and was able to enter the nucleus of meristematic cells, as suggested by several interactors localized in the nucleus, such as chromatin-associated factors. Finally, when proteins common to the two stages were considered, the two nuclear regulators OsNPR5/BOP1 and ARF12 appear to be strongly biotinylated under every condition by FTL1-TbID in rice meristems.

OsNPR5/BOP1 is an E3 ubiquitin ligase and transcriptional co-regulator. It contains a bric-a-brac/tramtrack/broad complex (BTB) domain as well as an ankyrin repeat, a very common protein-protein interaction domain found in bacteria and eukaryotes. The ankyrin repeat might function as substrate recognition domain to recruit targets that are selectively ubiquitinated and degraded by the 26S proteasome. Interestingly, we identified also the related NH5.1/OsBOP2 in stage 2 datasets. OsBOP1-3 are required to define the sheat:blade ratio of leaves, and triple loss-of-function mutants develop leaves with blade only (Toriba et al., 2019). Detailed temporal and spatial expression profiles indicate that *OsBOP* transcripts are localized in developing flowers, in all sterile organs, where FTL1 is also expressed (Giaume et al., 2023; Toriba et al., 2019). The molecular function of OsBOPs has not been defined yet. FTL1 could be a substrate targeted for degradation by the 26S proteasome. Removal of FTL1 protein in excess from the sterile organs of the flower, might be necessary to confer vegetative characteristics. Alternatively, FTL1 could be a co-regulator required for the activity of an OsBOP-containing complex. Interestingly, Arabidopsis BOP1 and BOP2 are required for limiting transcription of FD at the SAM, thereby reducing responsiveness of the meristem to floral induction mediated by FT (Andrés et al 2015). Double *bop1/bop2* mutants flower early under LD, whereas a dominant *BOP1-D* allele delays flowering and partially supresses the earliness of a *FT* overexpressor (Andrés et al., 2015). These data are consistent with a role in the photoperiodic pathway and part of this system might be conserved in rice. On the other hand, ARF12 takes part to several processes in rice biology, from phosphate homeostasis to plant immunity and vegetative development (Du et al., 2021; S. Wang et al., 2014; Z. Zhao et al., 2020). Its expression is regulated by OsmiR167 to determine male gametophyte formation and flower opening (H. Liu et al., 2012; Z.-X. Zhao et al., 2022) and is higher in roots and stems during the reproductive phase (RiceXPro). Its exact role as potential FTL1 interactor remains to be determined.

Localization predictions over the FTL1 interactome confirm the idea that FTL1, similarly to FT, could associate to the cell membrane, possibly through vesicle trafficking. This scenario is supported by literature describing a general propensity of plant PEBPs to be incorporated into the cell membrane or make use of components of the vesicle trafficking pathway (Jaillais & Parcy, 2021; Nakamura et al., 2014; Sohn et al., 2007; Susila et al., 2021, 2024; Zhu et al., 2021)(Jaillais & Parcy, 2021; Nakamura et al., 2014; Sohn et al., 2007; Susila et al., 2021, 2024; Zhu et al., 2021)(Jaillais & Parcy, 2021; Nakamura et al., 2014; Susila et al., 2021, 2024; Zhu et al., 2021)(Jaillais & Parcy, 2021; Nakamura et al., 2014; Susila et al., 2021, 2024; Zhu et al., 2021). Arabidopsis FT can localize to endomembranes, to the plasma membrane of phloem companion cells, and is loaded into the phloematic stream through the MCTP-SNARE endosomal trafficking system (L. Liu et al., 2019; Susila et al., 2021). Microtubule-associated and vesicle transport elements such as those enriched in our rice datasets are essential for proper cargo movement to the plasma membrane and/or out of it and indicate that also FTL1 could move within or between cells through vesicle-mediated transport. In rice, movement of Hd3a and RFT1 from companion cells to sieve elements is mediated by the MCTPs *Oryza sativa* FT INTERACTING PROTEIN 9 (OsFTIP9) and OsFTIP1, respectively (Song et al., 2017b; Zhang et al., 2022b). OsFTIP9, OsFTIP1, Hd3a and RFT1 co-localize with a marker of the endoplasmic reticulum. Therefore, proximity of FTL1 to components of the vesicle trafficking system might indicate similar mechanisms of transport.

Our experiments show that FTL1 is capable of moving to adjacent cells when expressed in single tobacco epidermal cells. The extent of movement was found to be similar to that of Hd3a and RFT1. This observation is not surprising, given the very high sequence identity between FTL1, Hd3a and RFT1, and their similar biochemical properties. However, whereas Hd3a and RFT1 are long-distance signals coming from the leaves, FTL1 protein is localized in all panicle meristems, with a pattern overlapping to its mRNAs, and suggestive of a cell-autonomous function. Cell-to-cell movement could occur locally in panicle meristems, raising questions related to the mechanisms of long-versus short-distance movement and its biological significance in rice. Movement between cells is very likely to be active because in our assays, a 2xGFP protein having size similar to FTL1-GFP could not move. Also, movement seems to be an intrinsic feature of FTL1, Hd3a and RFT1, because it occurred in a heterologous system without co-expression of other rice proteins, even if it cannot be excluded that tobacco proteins could have participated in cell-to-cell trafficking. Since movement was symplastic, we hypothesize a direct effect of FTL1, Hd3a and RFT1 on permeability of plasmodesmata. Proximity of FTL1 to components of vesicle trafficking might be related to movement between cells, as already mentioned. Similarly to Hd3a and RFT1, interaction with MCTPs could be involved. A recent study showed that Arabidopsis MCTPs act as tethers between the plasma membrane and the membrane of the desmotubule at membrane contact sites of plasmodesmata, and they are part of an active mechanism controlling cell-to-cell connectivity (Pérez-Sancho et al., 2025). We did not identify MCTPs by PL of FTL1. However, several lysines of MCTPs are required for interaction with the negatively charged phospholipids (phosphatidylinositol-4-phosphate in particular) of the plasma membrane and might not be accessible for biotinylation (Pérez-Sancho et al., 2025). Therefore, an MCTP-FTL1 interaction controlling plasmodesmata permeability is currently an attractive hypothesis to test.

In wider terms, our results remark the potentiality of PL for the study of tissue-specific dynamic processes and expand our understanding on the role of PEBP proteins in biological processes of agricultural relevance, such as rice panicle development.

## MATERIALS AND METHODS

### Plant growth conditions and sample preparation

The *Nicotiana benthamiana* plants used in this work were grown at 20°C and under short day (SD) conditions (8 h light - 16 h dark). Agroinfiltration was performed on 1- to 1.5-months-old green leaves at OD_600_=0,4. For PL preliminary tests and proteomic analysis, biotin infiltration and leaf sampling were carried out 3 days later.

Rice plants of the *O. sativa sb. japonica* var. Nipponbare were grown at an average temperature of 28°C under long day (LD) conditions (16 h light - 8 h dark). A shift to SD conditions (10 h light - 14 h dark) was employed to induce flowering in a controlled manner before sampling of reproductive meristems. For flowering time phenotyping under SD conditions, the photoperiodic shift was done 1 month after sowing.

Meristematic tissues indicative of sampling stages 1 and 2, as shown in Fig. 5, were sampled along with those for proteomic analyses, cleared with ethanol and observed under a Nikon Elicpse E600 Normasky microscope. Biological triplicates were employed for PL analyses. Each proteomic sample contained 30 and 15 flowering-induced meristems, for stage 2 and stage 1, respectively.

### Cloning of PL constructs and calli transformation

PL fusion constructs were obtained through the MultiSite Gateway system. AttB sequences were added by PCR to promoters and gene coding sequences for recombination into pDONOR221 P4-P1r and pDONOR221 P1-P2, respectively. Both TbID linker versions were synthesized by GeneWiz (https://www.genewiz.com) and recombined into pDONOR221 P2r-P3. Rice codon optimization was required according to the Codon Usage Database of Kazusa DNA Research Centre, Japan (https://www.kazusa.or.jp/codon). For each final construct, the three corresponding pENTRY clones were recombined into pDEST pH7m34GW through LR reaction. For the TbID construct, Gateway cloning was used for inserting the *TbID* sequence into the pB2GW7 destination vector.

The vectors were transformed into DB3.1 (chloramphenicol resistant) and DH5α *Escherichia coli* cells for the cloning steps, and into EHA105 A*grobacterium tumefaciens* cells for *N. benthamiana* agroinfiltrations and for the transformation of rice seed-derived calli. Calli induction, transformation and selection was executed as described in (Sahoo et al., 2011). The sequences of primers are reported in Supplementary Table 5.

### Protein extraction and precipitation

Nitrogen-frozen *N. benthamiana* and rice tissues were cold-grinded and lysed with a pH neutral buffer containing 4M Urea, 50mM Tris-HCl, 2mM EDTA, 150 mM NaCl, 0,2% 2-mercaptoethanol, 0,1 % SDS and Pierce Protease Inhibitors (Thermo Scientific™). The cell lysates were centrifuged at 14.000 rpm/4°C for 30 minutes, and the supernatant was recollected. Protein concentration was quantified using Quick Start Bradford 1x Dye Reagent (Bio-Rad™). Fresh extracts were normalized and filtered through Zeba Spin Desalting Columns, 7K MWCO (Thermo Scientific™) in pull down experiments. The filtered samples were incubated overnight at 4°C with Dynabeads MyOne Streptavidin C1 (Invitrogen™), to separate the biotinylated proteome. The beads were washed with a bland pH neutral buffer, containing 50 mM TrisHCl, 2 mM EDTA, 150 mM NaCl, 0,01% SDS and Pierce® Protease Inhibitors, and the pull downs were eluted in 2X Laemmli sample buffer.

### Western blot and silver staining

Protein samples were loaded onto a 10% acrylamide/bis-acrylamide gel (29:1 ratio). Electrophoresis was performed at a constant current of 30 mA for 1.5 hours. Before gel loading, raw protein extracts were combined with 4X Laemmli buffer. The separated proteins were transferred to a methanol-activated PVDF membrane by semi-dry blotting with Trans-Blot® Turbo™ Transfer System (1704150, Bio-Rad) after fast soaking into diluted Trans-Blot Turbo 5x Transfer Buffer (10026938, Bio-Rad). Protein transfer was performed at 45 mA for 7 minutes. For Western blotting, the membrane was incubated overnight at 4°C with the corresponding primary antibody. Following several washes, the membrane was incubated with the corresponding secondary antibody at RT for 1 hour. When Streptavidin, HRP conjugate (Millipore™, 18-152) was employed, one-step incubation was performed at RT for 2 hours. The membrane was developed using Clarity Max ECL substrate (Bio-Rad) and visualized using the ChemiDoc Imaging system (Bio-Rad). Silver staining was performed with Pierce^TM^ Silver Stain Kit (REF 24612) following the manufacturer’s instructions. The following antibodies were used in WB experiments: Anti-YFP mouse monoclonal, Agrisera™ (AS18 4177); Anti-RFP mouse monoclonal, Agrisera™ (AS15 3028); Anti-HA rabbit polyclonal, Agrisera™ (AS12 2220); Anti-mouse IgG, Bio-Rad™ (1706516); Anti-rabbit, Agrisera™ (AS09 602).

### Mass spectrometry and data analysis

The samples were analysed in biological triplicate for each experimental condition, using an Orbitrap Fusion Lumos ETD machine. Sample pre-processing LC-MS and data analyses have been performed by EMBL Proteomics Core Facility in Heidelberg-Germany 713 (https://www.embl.org/groups/proteomics), as described in Biancucci et al., (2025). Raw protein.tsv files from FragPipe were processed in R programming environment. Contaminants and reverse proteins were filtered out, and only proteins with ≥2 razor peptides were analyzed. Batch effects were corrected with the “removeBatchEffect” function from the limma package, and data were log2-transformed and normalized using the “normalizeVSN” function. Differential expression analysis was performed with limma’s moderated t-test. P-values and false discovery rates (FDRs) were calculated using the fdrtool function. Proteins with FDR<0.05 and an absolute fold change (FC) >2 were classified as hits, whereas those with FDR<0.2 and an absolute FC >1.5 were classified as candidates. Clustering of enriched proteins was performed using k-means with Euclidean distance and Ward.D2 linkage. The optimal number of clusters was as indicated by the Elbow method (4). Raw proteomics data are being uploaded on the PRIDE repository.

### Mobility Assays

Mobility assay experiments were conducted following the protocol described in Ohtsu et al. 2024. CDSs of the genes have been amplified from Nipponbare cDNA obtained from inflorescences. The reverse primers present an additional sequence encoding a 6-aminoacid linker (3XGly-Ser). The amplified sequences have been introduced in the FP08018 vector upon restriction with BsaI (ThermoFisher Scientific™) and ligation with T4 DNA ligase (Promega™). Final vectors have been introduced in *Agrobacterium tumefaciens* strain AGL1. The primers used are listed in Supplementary Table 5.

For the mobility assay experiment, constructs were infiltrated in tobacco leaves with an OD600 of 3×10^-5^; samples were observed at the confocal microscope 72h after the infiltration. The confocal microscope used was a Leica Stellaris 5 with LAS X software, together with the *TauGating* function, which was used for removing undesired signals derived from the autofluorescence of the samples. The excitation laser for the GFP was set at 489 nm, and emission spectra were collected in a 509 – 530 nm window. For dTomato, the excitation laser used was 561 nm, and emission was collected at 600-664 nm. Z-stack mode was used to acquire sequential plans, to capture all the cells in which the GFP had moved. ImageJ was used for processing the images. For mobility assays, statistical analyses have been performed with RStudio, using dyplr (v 1.1.4; https://dplyr.tidyverse.org/index.html), multcompView (v 0.1-10; Graves et al. 2024) and AICcmodavg (v 2.3-3; Mazerolle et al. 2023). Non-parametric statistical test Kruskal-Wallis was applied, with Dunn’s test as a post-hoc analysis. Results were summarized using a Compact Letter Display (CLD) to visualize statistically distinct means effectively.

## ACKNOWLEDGEMENTS

This work was supported by the University of Milan for PSR_Linea 3 FLORICE_My First SEED (MUR DM6 737/2021) awarded to CB and Agritech National Research Center and received funding from the European Union Next-GenerationEU (PIANO NAZIONALE DI RIPRESA E RESILIENZA (PNRR) – MISSIONE 4 COMPONENTE 2, INVESTIMENTO 1.4 – D.D. 1032 17/06/2022, CN00000022) for a postdoctoral fellowship to DC.

We are grateful to the personnel of the Botanical Garden ‘Città Studi’ for support with plant care, and to Jennifer Schwartz and Frank Stein of the EMBL Proteomics core facility for support with proteomic analyses. We thank the UNITECH NOLIMITS advanced microscopy facility of the University of Milan where imaging was carried out.

## References

Abelenda, J. A., Bergonzi, S., Oortwijn, M., Sonnewald, S., Du, M., Visser, R. G. F., Sonnewald, U., & Bachem, C. W. B. (2019). Source-sink regulation is mediated by interaction of an FT homolog with a SWEET protein in potato. Current Biology, 29(7), 1178–1186.

Alban, C. (2011). Biotin (vitamin B8) synthesis in plants. Advances in Botanical Research, 59, 39–66.

Alban, C., Job, D., & Douce, R. (2000). Biotin metabolism in plants. Annual Review of Plant Biology, 51(1), 17–47.

Andrés, F., Kinoshita, A., Kalluri, N., Fernández, V., Falavigna, V. S., Cruz, T. M. D., Jang, S., Chiba, Y., Seo, M., & Mettler-Altmann, T. (2020). The sugar transporter SWEET10 acts downstream of FLOWERING LOCUS T during floral transition of Arabidopsis thaliana. BMC Plant Biology, 20, 1–14.

Andrés, F., Romera-Branchat, M., Martínez-Gallegos, R., Patel, V., Schneeberger, K., Jang, S., Altmüller, J., Nürnberg, P., & Coupland, G. (2015). Floral induction in Arabidopsis by FLOWERING LOCUS T requires direct repression of BLADE-ON-PETIOLE genes by the homeodomain protein PENNYWISE. Plant Physiology, 169(3), 2187–2199.

Arora, D., Abel, N. B., Liu, C., Van Damme, P., Yperman, K., Eeckhout, D., Vu, L. D., Wang, J., Tornkvist, A., & Impens, F. (2020). Establishment of proximity-dependent biotinylation approaches in different plant model systems. Plant Cell, 32(11), 3388–3407.

Arul, A. B., & Robinson, R. A. S. (2018). Sample multiplexing strategies in quantitative proteomics. Analytical Chemistry, 91(1), 178–189.

Baldet, P., Alban, C., Axiotis, S., & Douce, R. (1993). Localization of free and bound biotin in cells from green pea leaves. Archives of Biochemistry and Biophysics, 303(1), 67–73.

Biancucci, M., Chirivì, D., Baldini, A., Badenhorst, E., Dobetti, F., Khahani, B., Formentin, E., Eguen, T., Turck, F., Moore, J. P., Tavakol, E., Wenkel, S., Lo Schiavo, F., Ezquer, I., Brambilla, V., Horner, D., Chiara, M., Perrella, G., Betti, C., & Fornara, F. (2025). Mutations in HEADING DATE 1 affect transcription and cell wall composition in rice. Plant Physiology, 197(4)10.1093/plphys/kiaf120

Branon, T. C., Bosch, J. A., Sanchez, A. D., Udeshi, N. D., Svinkina, T., Carr, S. A., Feldman, J. L., Perrimon, N., & Ting, A. Y. (2018). Efficient proximity labeling in living cells and organisms with TurboID. Nature Biotechnology, 36(9), 880–887.

Cerise, M., Giaume, F., Galli, M., Khahani, B., Lucas, J., Podico, F., Tavakol, E., Parcy, F., Gallavotti, A., Brambilla, V., & Fornara, F. (2021). OsFD4 promotes the rice floral transition via florigen activation complex formation in the shoot apical meristem. New Phytologist, 229(1), 429–443. 10.1111/nph.16834

Chaves-Sanjuan, A., Gnesutta, N., Gobbini, A., Martignago, D., Bernardini, A., Fornara, F., Mantovani, R., & Nardini, M. (2021). Structural determinants for NF-Y subunit organization and NF-Y/DNA association in plants. The Plant Journal, 105(1), 49–61. 10.1111/tpj.15038

Chirivì, D., & Betti, C. (2023). Molecular links between flowering and abiotic stress response: A focus on Poaceae. Plants, 12(2), 331.

Corbesier, L., Vincent, C., Jang, S., Fornara, F., Fan, Q., Searle, I., Giakountis, A., Farrona, S., Gissot, L., & Turnbull, C. (2007). FT protein movement contributes to long-distance signaling in floral induction of Arabidopsis. Science, 316(5827), 1030–1033.

Ding, C., Wang, Y., Chang, Z., You, S., Liu, Z., Wang, S., & Ding, Y. (2016). Comparative proteomic analysis reveals nitrogen fertilizer increases spikelet number per panicle in rice by repressing protein degradation and 14-3-3 proteins. Journal of Plant Growth Regulation, 35, 744–754.

Doi, K., Izawa, T., Fuse, T., Yamanouchi, U., Kubo, T., Shimatani, Z., Yano, M., & Yoshimura, A. (2004). Ehd1, a B-type response regulator in rice, confers short-day promotion of flowering and controls FT-like gene expression independently of Hd1. Genes & Development, 18(8), 926–936.

Du, X., Zhao, Y., Zhao, C., Zhao, S., Li, Y., Cheng, Y., Zhao, Q., Du, Y., Sun, H., & Sun, H. (2021). Construction of rice OsA RF12 *mutant based on CRISPR/Cas9 technology*.

Freytes, S. N., Canelo, M., & Cerdán, P. D. (2021). Regulation of flowering time: when and where? Current Opinion in Plant Biology, 63, 102049.

Fridy, P. C., Thompson, M. K., Ketaren, N. E., & Rout, M. P. (2015). Engineered high-affinity nanobodies recognizing staphylococcal Protein A and suitable for native isolation of protein complexes. Analytical Biochemistry, 477, 92– 94.

Giaume, F., Bono, G. A., Martignago, D., Miao, Y., Vicentini, G., Toriba, T., Wang, R., Kong, D., Cerise, M., & Chirivì, D. (2023). Two florigens and a florigen-like protein form a triple regulatory module at the shoot apical meristem to promote reproductive transitions in rice. Nature Plants, 9(4), 525–534.

Gómez-Ariza, J., Brambilla, V., Vicentini, G., Landini, M., Cerise, M., Carrera, E., Shrestha, R., Chiozzotto, R., Galbiati, F., & Caporali, E. (2019). A transcription factor coordinating internode elongation and photoperiodic signals in rice. Nature Plants, 5(4), 358–362.

Goretti, D., Martignago, D., Landini, M., Brambilla, V., Gómez-Ariza, J., Gnesutta, N., Galbiati, F., Collani, S., Takagi, H., Terauchi, R., Mantovani, R., & Fornara, F. (2017). Transcriptional and Post-transcriptional Mechanisms Limit Heading Date 1 (Hd1) Function to Adapt Rice to High Latitudes. PLOS Genetics, 13(1), e1006530-. 10.1371/journal.pgen.1006530

Grismer, T. S., Karundasa, S. S., Shrestha, R., Byun, D., Ni, W., Reyes, A. V, & Xu, S.-L. (2024). Workflow enhancement of TurboID-mediated proximity labeling for SPY signaling network mapping. BioRxiv.

Guo, J., Guo, S., Lu, S., Gong, J., Wang, L., Ding, L., Chen, Q., & Liu, W. (2023). The development of proximity labeling technology and its applications in mammals, plants, and microorganisms. Cell Communication and Signaling, 21(1), 269. 10.1186/s12964-023-01310-1

Ho, W. W. H., & Weigel, D. (2014). Structural features determining flower-promoting activity of Arabidopsis FLOWERING LOCUS T. The Plant Cell, 26(2), 552–564.

Hou, S., Shi, L., & Lei, H. (2016). Biotin–Streptavidin Affinity Purification of RNA–Protein Complexes Assembled In Vitro. RNA-Protein Complexes and Interactions: Methods and Protocols, 23–34.

Itoh, H., Nonoue, Y., Yano, M., & Izawa, T. (2010). A pair of floral regulators sets critical day length for Hd3a florigen expression in rice. Nature Genetics, 42(7), 635–638.

Jaillais, Y., & Parcy, F. (2021). Lipid-mediated regulation of flowering time. Science, 373(6559), 1086–1087.

Jang, S., Li, H.-Y., & Kuo, M.-L. (2017). Ectopic expression of Arabidopsis FD and FD PARALOGUE in rice results in dwarfism with size reduction of spikelets. Scientific Reports, 7(1), 44477.

Kaneko-Suzuki, M., Kurihara-Ishikawa, R., Okushita-Terakawa, C., Kojima, C., Nagano-Fujiwara, M., Ohki, I., Tsuji, H., Shimamoto, K., & Taoka, K.-I. (2018). TFL1-like proteins in rice antagonize rice FT-like protein in inflorescence development by competition for complex formation with 14-3-3 and FD. Plant and Cell Physiology, 59(3), 458–468.

Kang, M.-G., & Rhee, H.-W. (2022). Molecular spatiomics by proximity labeling. Accounts of Chemical Research, 55(10), 1411–1422.

Kaur, A., Nijhawan, A., Yadav, M., & Khurana, J. P. (2021). OsbZIP62/OsFD7, a functional ortholog of FLOWERING LOCUS D, regulates floral transition and panicle development in rice. Journal of Experimental Botany, 72(22), 7826–7845.

Kean-Galeno, T., Lopez-Arredondo, D., & Herrera-Estrella, L. (2024). The Shoot Apical Meristem: An Evolutionary Molding of Higher Plants. International Journal of Molecular Sciences, 25(3), 1519.

Kim, D. I., & Roux, K. J. (2016). Filling the void: proximity-based labeling of proteins in living cells. Trends in Cell Biology, 26(11), 804–817.

Kim, H. B., & Kim, K. (2024). Precision proteomics with TurboID: mapping the suborganelle landscape. The Korean Journal of Physiology & Pharmacology: Official Journal of the Korean Physiological Society and the Korean Society of Pharmacology, 28(6), 495–501.

Kim, T.-W., Park, C. H., Hsu, C.-C., Kim, Y.-W., Ko, Y.-W., Zhang, Z., Zhu, J.-Y., Hsiao, Y.-C., Branon, T., & Kaasik, K. (2023). Mapping the signaling network of BIN2 kinase using TurboID-mediated biotin labeling and phosphoproteomics. The Plant Cell, 35(3), 975–993.

Komiya, R., Ikegami, A., Tamaki, S., Yokoi, S., & Shimamoto, K. (2008). Hd3a and RFT1 are essential for flowering in rice.

Lauschke, A., Maibaum, L., Engel, M., Eisengräber, L., Bayer, S., Hackel, A., & Kühn, C. (2025). The potato sugar transporter SWEET1g affects apoplasmic sugar ratio and phloem-mobile tuber-and flower-inducing signals. Plant Physiology, 197(1), kiae602.

Lee, M.-H., Jackson, M. A., Rehm, F. B. H., Barkauskas, D. S., Ho, W. L., Yap, K., Craik, D. J., & Gilding, E. K. (2024). Proximity Labelling Confirms the Involvement of Papain-Like Cysteine Proteases and Chaperones in Cyclotide Biosynthesis. Plant Molecular Biology Reporter, 1–13.

Li, G., Zhang, H., Li, J., Zhang, Z., & Li, Z. (2021). Genetic control of panicle architecture in rice. The Crop Journal, 9(3), 590–597

Li, P., Li, J., Wang, L., & Di, L. (2017). Proximity labeling of interacting proteins: application of BioID as a discovery tool. Proteomics, 17(20), 1700002.

Li, X., Wei, Y., Fei, Q., Fu, G., Gan, Y., & Shi, C. (2023). TurboID-mediated proximity labeling for screening interacting proteins of FIP37 in Arabidopsis. Plant Direct, 7(12), e555.

Li, Z., Liu, S.-L., Montes-Serey, C., Walley, J. W., & Aung, K. (2022). Plasmodesmata-located proteins regulate plasmodesmal function at specific cell interfaces in Arabidopsis. BioRxiv, 2022–2028.

Lin, Q., Zhou, Z., Luo, W., Fang, M., Li, M., & Li, H. (2017). Screening of proximal and interacting proteins in rice protoplasts by proximity-dependent biotinylation. Frontiers in Plant Science, 8, 749.

Lin, Z., Schaefer, K., Lui, I., Yao, Z., Fossati, A., Swaney, D. L., Palar, A., Sali, A., & Wells, J. A. (2024). Multiscale photocatalytic proximity labeling reveals cell surface neighbors on and between cells. Science, 385(6706), eadl5763.

Liu, H., Jia, S., Shen, D., Liu, J., Li, J., Zhao, H., Han, S., & Wang, Y. (2012). Four AUXIN RESPONSE FACTOR genes downregulated by microRNA167 are associated with growth and development in Oryza sativa. Functional Plant Biology, 39(9), 736–744.

Liu, L., Li, C., Teo, Z. W. N., Zhang, B., & Yu, H. (2019). The MCTP-SNARE Complex Regulates Florigen Transport in Arabidopsis. The Plant Cell, 31(10), 2475–2490. 10.1105/tpc.18.00960

Macknight, R., Bancroft, I., Page, T., Lister, C., Schmidt, R., Love, K., Westphal, L., Murphy, G., Sherson, S., Cobbett, C., & Dean, C. (1997). FCA, a Gene Controlling Flowering Time in Arabidopsis, Encodes a Protein Containing RNA-Binding Domains. Cell, 89(5), 737–745. 10.1016/S0092-8674(00)80256-1

Mair, A., Xu, S.-L., Branon, T. C., Ting, A. Y., & Bergmann, D. C. (2019). Proximity labeling of protein complexes and cell-type-specific organellar proteomes in Arabidopsis enabled by TurboID. Elife, 8, e47864.

May, D. G., Scott, K. L., Campos, A. R., & Roux, K. J. (2020). Comparative application of BioID and TurboID for protein-proximity biotinylation. Cells, 9(5), 1070.

Mineri, L., Bono, G. A., Sergi, E., Colleoni, P. E., Morandini, P., Vicentini, G., Fornara, F., & Brambilla, V. (2025). OsMAINTENANCE OF MERISTEM LIKE 1 controls style number at high temperatures in rice. Plant Molecular Biology, 115(1), 24.

Mineri, L., Cerise, M., Giaume, F., Vicentini, G., Martignago, D., Chiara, M., Galbiati, F., Spada, A., Horner, D., & Fornara, F. (2023). Rice florigens control a common set of genes at the shoot apical meristem including the F- BOX BROADER TILLER ANGLE 1 that regulates tiller angle and spikelet development. The Plant Journal, 115(6), 1647–1660.

Mishra, S. K., Tripp, J., Winkelhaus, S., Tschiersch, B., Theres, K., Nover, L., & Scharf, K.-D. (2002). In the complex family of heat stress transcription factors, HSfA1 has a unique role as master regulator of thermotolerance in tomato. Genes & Development, 16, 1555–1567. 10.1101/gad.228802

Nakamura, Y., Andrés, F., Kanehara, K., Liu, Y., Dörmann, P., & Coupland, G. (2014). Arabidopsis florigen FT binds to diurnally oscillating phospholipids that accelerate flowering. Nature Communications, 5(1), 3553. 10.1038/ncomms4553

Navarro, C., Abelenda, J. A., Cruz-Oró, E., Cuéllar, C. A., Tamaki, S., Silva, J., Shimamoto, K., & Prat, S. (2011). Control of flowering and storage organ formation in potato by FLOWERING LOCUS T. Nature, 478(7367), 119– 122.

Ohtsu, M., Jennings, J., Johnston, M., Breakspear, A., Liu, X., Stark, K., Morris, R. J., de Keijzer, J., & Faulkner, C. (2024). Assaying effector cell-to-cell mobility in plant tissues identifies hypermobility and indirect manipulation of plasmodesmata. Molecular Plant-Microbe Interactions, 37(2), 84–92.

Pappireddi, N., Martin, L., & Wühr, M. (2019). A review on quantitative multiplexed proteomics. Chembiochem, 20(10), 1210–1224.

Pérez-Sancho, J., Smokvarska, M., Dubois, G., Glavier, M., Sritharan, S., Moraes, T. S., Moreau, H., Dietrich, V., Platre, M. P., & Paterlini, A. (2025). Plasmodesmata act as unconventional membrane contact sites regulating intercellular molecular exchange in plants. Cell, 188(4), 958–977.

Purwestri, Y. A., Ogaki, Y., Tamaki, S., Tsuji, H., & Shimamoto, K. (2009). The 14-3-3 protein GF14c acts as a negative regulator of flowering in rice by interacting with the florigen Hd3a. Plant and Cell Physiology, 50(3), 429– 438.

Qin, W., Cho, K. F., Cavanagh, P. E., & Ting, A. Y. (2021). Deciphering molecular interactions by proximity labeling. Nature Methods, 18(2), 133–143.

Rakk, D., Kukolya, J., Škrbić, B. D., Vágvölgyi, C., Varga, M., & Szekeres, A. (2023). Advantages of multiplexing ability of the orbitrap mass analyzer in the multi-mycotoxin analysis. Toxins, 15(2), 134.

Roux, K. J., Kim, D. I., Raida, M., & Burke, B. (2012). A promiscuous biotin ligase fusion protein identifies proximal and interacting proteins in mammalian cells. Journal of Cell Biology, 196(6), 801–810.

Sahoo, K. K., Tripathi, A. K., Pareek, A., Sopory, S. K., & Singla-Pareek, S. L. (2011). An improved protocol for efficient transformation and regeneration of diverse indica rice cultivars. Plant Methods, 7(1), 49. 10.1186/1746-4811-7-49

Samavarchi-Tehrani, P., Samson, R., & Gingras, A.-C. (2020). Proximity dependent biotinylation: key enzymes and adaptation to proteomics approaches. Molecular & Cellular Proteomics, 19(5), 757–773.

Serre, L., Vallée, B., Bureaud, N., Schoentgen, F., & Zelwer, C. (1998). Crystal structure of the phosphatidylethanolamine-binding protein from bovine brain: a novel structural class of phospholipid-binding proteins. Structure, 6(10), 1255–1265.

Shen, C., Liu, H., Guan, Z., Yan, J., Zheng, T., Yan, W., Wu, C., Zhang, Q., Yin, P., & Xing, Y. (2020). Structural Insight into DNA Recognition by CCT/NF-YB/YC Complexes in Plant Photoperiodic Flowering. The Plant Cell, 32(11), 3469–3484. 10.1105/tpc.20.00067

Shi, L., Ferrando, T. M., Villanueva, S. L., Joosten, M. H. A. J., Vleeshouwers, V. G. A. A., & Bachem, C. W. B. (2023a). Protocol to identify protein-protein interaction networks in Solanum tuberosum using transient TurboID-based proximity labeling. STAR Protocols, 4(4), 102577.

Shi, L., Ferrando, T. M., Villanueva, S. L., Joosten, M. H. A. J., Vleeshouwers, V. G. A. A., & Bachem, C. W. B. (2023b). Protocol to identify protein-protein interaction networks in Solanum tuberosum using transient TurboID-based proximity labeling. STAR Protocols, 4(4), 102577.

Sohn, E. J., Rojas-Pierce, M., Pan, S., Carter, C., Serrano-Mislata, A., Madueño, F., Rojo, E., Surpin, M., & Raikhel, N. V. (2007). The shoot meristem identity gene TFL1 is involved in flower development and trafficking to the protein storage vacuole. Proceedings of the National Academy of Sciences, 104(47), 18801–18806. 10.1073/pnas.0708236104

Song, S., Chen, Y., Liu, L., Wang, Y., Bao, S., Zhou, X., Teo, Z. W. N., Mao, C., Gan, Y., & Yu, H. (2017a). OsFTIP1-mediated regulation of florigen transport in rice is negatively regulated by the ubiquitin-like domain kinase OsUbDKγ4. The Plant Cell, 29(3), 491–507.

Song, S., Chen, Y., Liu, L., Wang, Y., Bao, S., Zhou, X., Teo, Z. W. N., Mao, C., Gan, Y., & Yu, H. (2017b). OsFTIP1-mediated regulation of florigen transport in rice is negatively regulated by the ubiquitin-like domain kinase OsUbDKγ4. The Plant Cell, 29(3), 491–507.

Susila, H., Gawarecka, K., Youn, G., Jurić, S., Jeong, H., & Ahn, J. H. (2024). THYLAKOID FORMATION 1 interacts with FLOWERING LOCUS T and modulates temperature-responsive flowering in Arabidopsis. The Plant Journal, 120(1), 60–75.

Susila, H., Jurić, S., Liu, L., Gawarecka, K., Chung, K. S., Jin, S., Kim, S.-J., Nasim, Z., Youn, G., Suh, M. C., Yu, H., & Ahn, J. H. (2021). Florigen sequestration in cellular membranes modulates temperature-responsive flowering. Science, 373(6559), 1137–1142. 10.1126/science.abh4054

Tamaki, S., Tsuji, H., Matsumoto, A., Fujita, A., Shimatani, Z., Terada, R., Sakamoto, T., Kurata, T., & Shimamoto, K. (2015). FT-like proteins induce transposon silencing in the shoot apex during floral induction in rice. Proceedings of the National Academy of Sciences, 112(8), E901–E910.

Tan, H., Zhou, Y., Dinius, E., & Lozano-Durán, R. (2024). The Ti-TAN plasmid toolbox for TurboID-based proximity labeling assays in Nicotiana benthamiana. Journal of Integrative Plant Biology, 66(2), 166–168.

Tan, W., Chen, J., Yue, X., Chai, S., Liu, W., Li, C., Yang, F., Gao, Y., Gutiérrez Rodríguez, L., Resco de Dios, V., Zhang,D., & Yao, Y. (2025). The heat response regulators HSFA1s promote Arabidopsis thermomorphogenesis via stabilizing PIF4 during the day. Science Advances, 9(44), eadh1738. 10.1126/sciadv.adh1738

Tang, Y., Yang, X., Huang, A., Seong, K., Ye, M., Li, M., Zhao, Q., Krasileva, K., & Gu, Y. (2024). Proxiome assembly of the plant nuclear pore reveals an essential hub for gene expression regulation. Nature Plants, 1–13.

Taoka, K., Ohki, I., Tsuji, H., Furuita, K., Hayashi, K., Yanase, T., Yamaguchi, M., Nakashima, C., Purwestri, Y. A., & Tamaki, S. (2011). 14-3-3 proteins act as intracellular receptors for rice Hd3a florigen. Nature, 476(7360), 332–335.

Toriba, T., Tokunaga, H., Shiga, T., Nie, F., Naramoto, S., Honda, E., Tanaka, K., Taji, T., Itoh, J.-I., & Kyozuka, J. (2019). BLADE-ON-PETIOLE genes temporally and developmentally regulate the sheath to blade ratio of rice leaves. Nature Communications, 10(1), 619. 10.1038/s41467-019-08479-5

Toribio, R., Navarro, A., & Castellano, M. M. (2024). HOP stabilizes the HSFA1a and plays a main role in the onset of thermomorphogenesis. Plant, Cell & Environment, 47(11), 4449–4463. 10.1111/pce.15036

Tsuji, H., Nakamura, H., Taoka, K., & Shimamoto, K. (2013). Functional diversification of FD transcription factors in rice, components of florigen activation complexes. Plant and Cell Physiology, 54(3), 385–397.

Tsuji, H., & Sato, M. (2024). The function of florigen in the vegetative-to-reproductive phase transition in and around the shoot apical meristem. Plant and Cell Physiology, 65(3), 322–337.

Tsuji, H., Tachibana, C., Tamaki, S., Taoka, K., Kyozuka, J., & Shimamoto, K. (2015). Hd3a promotes lateral branching in rice. The Plant Journal, 82(2), 256–266.

Vicentini, G., Biancucci, M., Mineri, L., Chirivì, D., Giaume, F., Miao, Y., Kyozuka, J., Brambilla, V., Betti, C., & Fornara, F. (2023). Environmental control of rice flowering time. Plant Communications, 4(5).

Wallner, E.-S., Mair, A., Handler, D., McWhite, C., Xu, S.-L., Dolan, L., & Bergmann, D. C. (2024). Spatially resolved proteomics of the Arabidopsis stomatal lineage identifies polarity complexes for cell divisions and stomatal pores. Developmental Cell, 59(9), 1096–1109.

Wang, S., Zhang, S., Sun, C., Xu, Y., Chen, Y., Yu, C., Qian, Q., Jiang, D., & Qi, Y. (2014). Auxin response factor (Os ARF 12), a novel regulator for phosphate homeostasis in rice (Oryza sativa). New Phytologist, 201(1), 91– 103.

Wang, Y.-C., & Chen, B.-S. (2010). Integrated cellular network of transcription regulations and protein-protein interactions. BMC Systems Biology, 4(1), 20. 10.1186/1752-0509-4-20

Xiong, Z., Lo, H. P., McMahon, K.-A., Parton, R. G., & Hall, T. E. (2021). Proximity dependent biotin labelling in zebrafish for Proteome and Interactome Profiling. Bio-Protocol, 11(19), e4178–e4178.

Xu, Y., Fan, X., & Hu, Y. (2021). In vivo interactome profiling by enzyme-catalyzed proximity labeling. Cell & Bioscience, 11(1), 27.

Yan, Z., Liang, D., Liu, H., & Zheng, G. (2010). FLC: A key regulator of flowering time in Arabidopsis. Russian Journal of Plant Physiology, 57, 166– 174.

Yano, M., Katayose, Y., Ashikari, M., Yamanouchi, U., Monna, L., Fuse, T., Baba, T., Yamamoto, K., Umehara, Y., & Nagamura, Y. (2000). Hd1, a major photoperiod sensitivity quantitative trait locus in rice, is closely related to the Arabidopsis flowering time gene CONSTANS. The Plant Cell, 12(12), 2473– 2483.

Zhang, L., Zhang, F., Zhou, X., Poh, T. X., Xie, L., Shen, J., Yang, L., Song, S., Yu, H., & Chen, Y. (2022a). The tetratricopeptide repeat protein OsTPR075 promotes heading by regulating florigen transport in rice. The Plant Cell, 34(10), 3632–3646.

Zhang, L., Zhang, F., Zhou, X., Poh, T. X., Xie, L., Shen, J., Yang, L., Song, S., Yu, H., & Chen, Y. (2022b). The tetratricopeptide repeat protein OsTPR075 promotes heading by regulating florigen transport in rice. The Plant Cell, 34(10), 3632–3646.

Zhao, Z., Feng, Q., Cao, X., Zhu, Y., Wang, H., Chandran, V., Fan, J., Zhao, J., Pu, M., & Li, Y. (2020). Osa-miR167d facilitates infection of Magnaporthe oryzae in rice. Journal of Integrative Plant Biology, 62(5), 702–715.

Zhao, Z.-X., Yin, X.-X., Li, S., Peng, Y.-T., Yan, X.-L., Chen, C., Hassan, B., Zhou, S.-X., Pu, M., & Zhao, J.-H. (2022). miR167d-ARF s Module Regulates Flower Opening and Stigma Size in Rice. Rice, 15(1), 40.

Zheng, T., Sun, J., Zhou, S., Chen, S., Lu, J., Cui, S., Tian, Y., Zhang, H., Cai, M., Zhu, S., Wu, M., Wang, Y., Jiang, L., Zhai, H., Wang, H., & Wan, J. (2019). Post-transcriptional regulation of Ghd7 protein stability by phytochrome and OsGI in photoperiodic control of flowering in rice. New Phytologist, 224(1), 306–320. 10.1111/nph.16010

Zhu, Y., Klasfeld, S., Jeong, C. W., Jin, R., Goto, K., Yamaguchi, N., & Wagner, D. (2020). TERMINAL FLOWER 1-FD complex target genes and competition with FLOWERING LOCUS T. Nature Communications, 11(1), 5118.

Zhu, Y., Klasfeld, S., & Wagner, D. (2021). Molecular regulation of plant developmental transitions and plant architecture via PEPB family proteins: an update on mechanism of action. Journal of Experimental Botany, 72(7), 2301–2311. 10.1093/jxb/eraa598

