## Supplementary Figures and tables for "The proximal proteome of FLOWERING LOCUS T LIKE 1 during rice panicle development suggests cell-to-cell mobility features": Supplementary Figures.pptx

### Slide 1
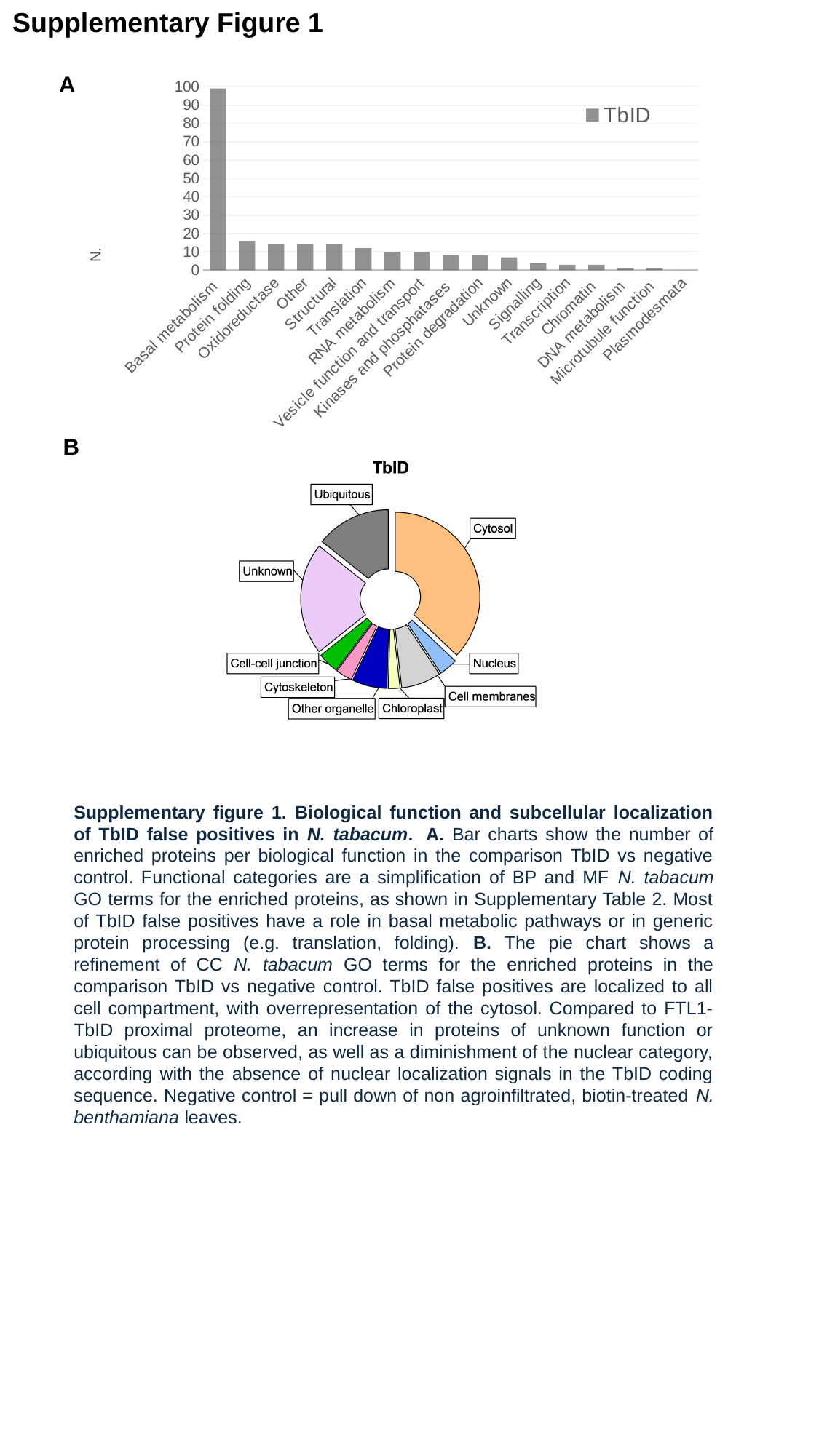

Supplementary Figure 1
A
#### Chart
| Category | TbID |
|---|---|
| Basal metabolism | 99.0 |
| Protein folding | 16.0 |
| Oxidoreductase | 14.0 |
| Other | 14.0 |
| Structural | 14.0 |
| Translation | 12.0 |
| RNA metabolism | 10.0 |
| Vesicle function and transport | 10.0 |
| Kinases and phosphatases | 8.0 |
| Protein degradation | 8.0 |
| Unknown | 7.0 |
| Signalling | 4.0 |
| Transcription | 3.0 |
| Chromatin | 3.0 |
| DNA metabolism | 1.0 |
| Microtubule function | 1.0 |
| Plasmodesmata | 0.0 |B
Supplementary figure 1. Biological function and subcellular localization of TbID false positives in N. tabacum.  A. Bar charts show the number of enriched proteins per biological function in the comparison TbID vs negative control. Functional categories are a simplification of BP and MF N. tabacum GO terms for the enriched proteins, as shown in Supplementary Table 2. Most of TbID false positives have a role in basal metabolic pathways or in generic protein processing (e.g. translation, folding). B. The pie chart shows a refinement of CC N. tabacum GO terms for the enriched proteins in the comparison TbID vs negative control. TbID false positives are localized to all cell compartment, with overrepresentation of the cytosol. Compared to FTL1-TbID proximal proteome, an increase in proteins of unknown function or ubiquitous can be observed, as well as a diminishment of the nuclear category, according with the absence of nuclear localization signals in the TbID coding sequence. Negative control = pull down of non agroinfiltrated, biotin-treated N. benthamiana leaves.

### Slide 2
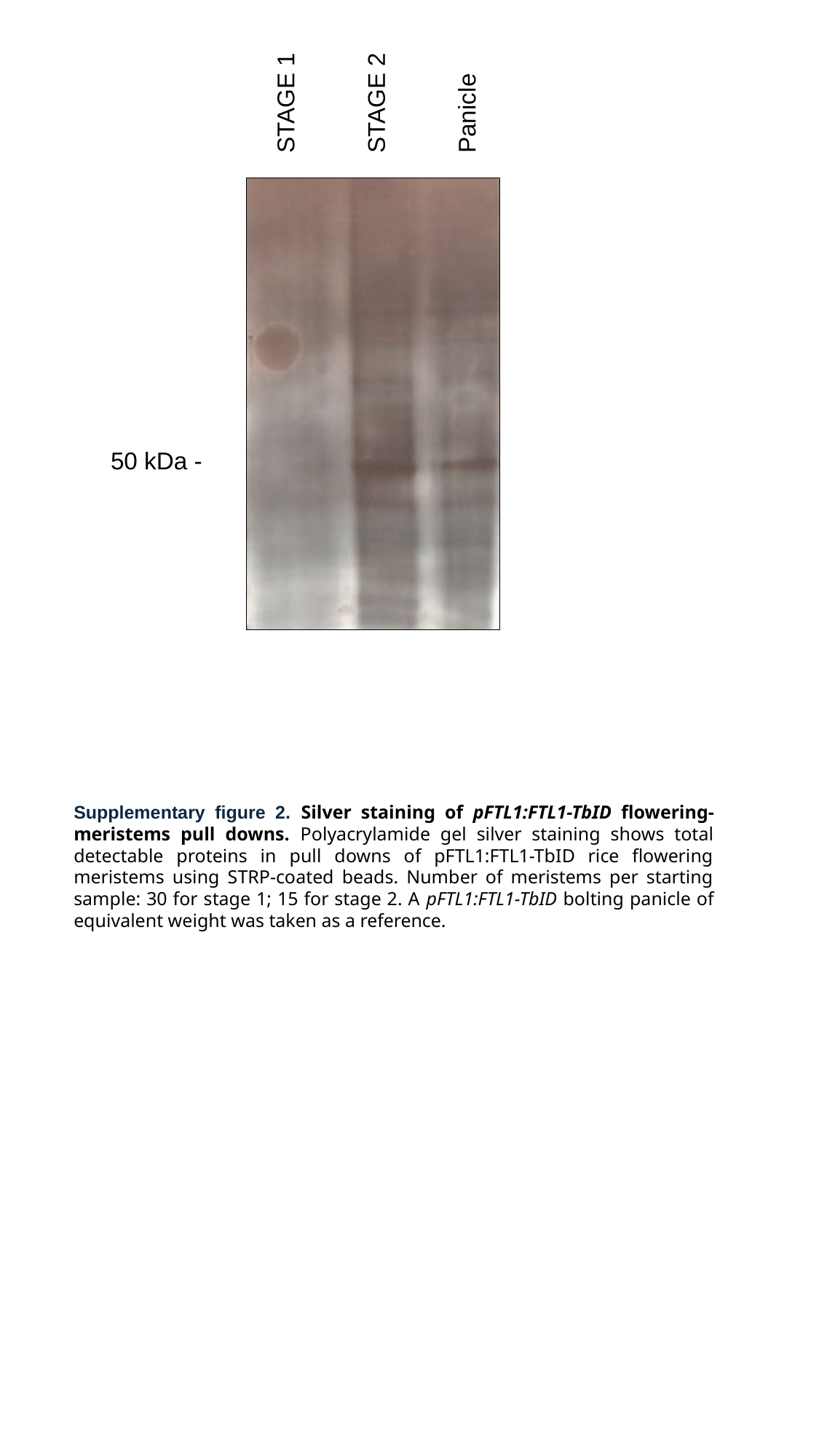

STAGE 2
STAGE 1
Panicle
50 kDa -
Supplementary figure 2. Silver staining of pFTL1:FTL1-TbID flowering-meristems pull downs. Polyacrylamide gel silver staining shows total detectable proteins in pull downs of pFTL1:FTL1-TbID rice flowering meristems using STRP-coated beads. Number of meristems per starting sample: 30 for stage 1; 15 for stage 2. A pFTL1:FTL1-TbID bolting panicle of equivalent weight was taken as a reference.
