## Supplementary Figures and tables for "The proximal proteome of FLOWERING LOCUS T LIKE 1 during rice panicle development suggests cell-to-cell mobility features": Supplementary Table 5.docx

**Supplementary table 5**. List of primers used in this study.

| **Primer name** | **Primer sequence** |
| --- | --- |
| CaMV p35S (Fw) + AttB4 | GGGGACAACTTTGTATAGAAAAGTTGCCTGAGACTTTTCAACAAAGGGT |
| CaMV p35S (Rv) + AttB1r | GGGGACTGCTTTTTTGTACAAACTTGCGTCCTCTCCAAATGAAATGAAC |
| FTL1 (Fw) + AttB1 | GGGGACAAGTTTGTACAAAAAAGCAGGCTTAATGAGCGGGCGGGGGAGG |
| FTL1 (Rv) without stop codon + AttB2 | GGGGACCACTTTGTACAAGAAAGCTGGGTGCATTCTTCTTCCTCCGGTT |
| pFTL1 (Fw) + AttB4 | GGGGACAACTTTGTATAGAAAAGTTGCCGTTTTGTTGCCCGATTCCCT |
| pFTL1 (Rv) + AttB1r | GGGGACTGCTTTTTTGTACAAACTTGCCACCACCACCCAGCCTTATA |
| TurboID (Fw) + AttB1 | GGGGACAAGTTTGTACAAAAAAGCAGGCTCTATGAAGGACAACACCGTGCC |
| 3xHA TAG (Rv) with stop codon + AttB2 | GGGGACCACTTTGTACAAGAAAGCTGGGTCTCAGGCGTAGTCCGGCAC |
| Genotyping TbID (Fw) | GcACcAAcCAgTAcCTccTc |
| Genotyping TbID (Rv) | TCcTTGTCgCCgATGATgAg |
| Genotyping FLT1 (Fw) | CTACACCCTGGTGATGGTGG |
| Genotyping pFLT1 (Fw) | TAGTTGTCCGGCCGATTCTT |
| Mobility FTL1 + BsaI restriction site (Fw) | CTATCTGGTCTCAAATGATGAGCGGGCGGGGGAGG |
| Mobility FTL1 + BsaI restriction site (Rv) | ATGCATGGTCTCTCGAACCCGATCCCGATCCCATTCTTCTTCCTCCGGTTC |
| Mobility Hd3a + BsaI restriction site (Fw) | CTATCTGGTCTCAAATGATGGCCGGAAGTGGCAGGGA |
| Mobility Hd3a + BsaI restriction site (Rv) | ATGCATGGTCTCTCGAACCCGATCCCGATCCGGGGTAGACCCTCCTGCCGC |
| Mobility RFT1 + BsaI restriction site (Fw) | CTATCTGGTCTCAAATGATGGCCGGCAGCGGCAGGGA |
| Mobility RFT1 + BsaI restriction site (Rv) | ATGCATGGTCTCTCGAACCCGATCCCGATCCGGGGTAGACCCTCCTGCCGC |
